# Distinct phenotypic consequences of cholangiocarcinoma-associated FGFR2 alterations depend on biliary epithelial cell state

**DOI:** 10.1101/2024.08.30.610360

**Authors:** C Chiasson-MacKenzie, Y Zhang, E O’Loughlin, PS Dave, AS Menon, V Vijay, AD Westerfield, V Kumar, SN Bhatia, E Rheinbay, N Bardeesy, SL Stott, AI McClatchey

**Author notes:** Equal contribution (C Chiasson-MacKenzie, Y Zhang, E O’Loughlin).

## Abstract

Epithelial cancers disrupt tissue architecture and are often driven by mutations in genes that play important roles in normal epithelial morphogenesis. The intrahepatic biliary system is an epithelial tubular network that forms within the developing liver via the *de novo* initiation and expansion of apical lumens. Intrahepatic biliary tumors (intrahepatic cholangiocarcinoma) commonly harbor activating genomic alterations in the FGFR2 receptor tyrosine kinase, which plays important roles in epithelial morphogenesis in other developmental settings. Using a physiologic and quantitative 3D model we demonstrate that FGFR signaling is important for biliary morphogenesis and that oncogenic FGFR2 fusions and in-frame deletions disrupt biliary architecture. Importantly, we show that the trafficking of and signaling from the FGFR2 mutants, as well as their phenotypic impacts, are governed by the epithelial state of the cell. Unexpectedly, we also found that distinct tumor-driving FGFR2 mutants disrupt biliary morphogenesis in completely different and clinically relevant ways, informing our understanding of morphogenesis and tumorigenesis and highlighting the importance of convergent studies of both.

**Summary statement:** Using a physiologic 3D model of biliary tubulogenesis, Chiasson-MacKenzie et al. show that FGFR2 signaling is important for biliary morphogenesis and that different intrahepatic cholangiocarcinoma-causing FGFR2 alterations have unexpectedly distinct and epithelial state- specific impacts on biliary architecture.

## Introduction

The acquisition of epithelial polarity and architecture is a cellular program that governs the morphogenesis of many organs, and abnormalities in epithelial tissue architecture are the first signs of cancer pathology triggered by somatic genetic mutations^1–4^. Despite the clear relationship between morphogenesis and tumorigenesis, tumor-causing genomic alterations are usually studied in cells cultured under conditions that do not recapitulate epithelial architecture, with changes in cell number used as the sole readout of activity. Alterations in receptor tyrosine kinases (RTKs) are common tumor drivers that are generally assumed to confer broad and constitutive signaling and unscheduled cell division^5^. However, RTK activity also governs and is modulated by programs of epithelial polarization and maturation^6,7^. In addition, germline activating mutations in certain RTKs cause specific developmental syndromes without increased cancer predisposition, while somatic genomic activation of RTKs is strongly biased across different cancer types. These patterns suggest important, but poorly understood biological relationships between RTKs and tumor context^8^. Understanding these relationships will require convergent studies of morphogenesis and tumorigenesis.

The epithelial lineages of the mature liver – hepatocytes and cholangiocytes (intrahepatic bile duct [IHBD] cells) – develop from common embryonic progenitors known as hepatoblasts. IHBDs form a tubular tree within the liver that is essential for liver function^9^. Biliary morphogenesis occurs in an ‘inside-out’ manner, driven by the *de novo* formation, extension and interconnection of apical lumens within E-cadherin-positive cell-cell junctions, and is accompanied by the conversion of immature hepatoblasts to a biliary fate^10,11^. This fundamental self-organizing mechanism of epithelial morphogenesis provides spatial control of both secretory and receptor activity in other developmental settings^12–14^. Bile duct formation is thus driven by the cellular acquisition of epithelial polarity, but little is known about how this program is initiated or coordinated among cells to establish a biliary tree.

Congenital or acquired diseases that are associated with abnormal architecture of the intrahepatic biliary epithelial tree are largely fatal, with intrahepatic cholangiocarcinoma representing a particularly aggressive malignancy in this context^15,16^. Genomic alterations in the fibroblast growth factor receptor 2 (FGFR2), are common in intrahepatic cholangiocarcinoma (ICC) and predict clinical response to pharmacologic FGFR inhibitors although acquired resistance limits overall efficacy^17–20^. In established ICC, FGFR2-driven signaling has been shown to sustain tumor growth by MAPK-activation and downstream transcriptional programs. By contrast, the role of FGFR2 in tumor initiation has been far less studied. In other developmental contexts, FGFR signaling is essential for the acquisition of epithelial architecture and, in particular, for *de novo* lumen formation^13,14,21,22^. Yet, the biological function of FGFR in biliary morphogenesis as well as the proximal consequences of its unscheduled activation in biliary tumorigenesis remain unknown.

Much has been learned about biliary morphogenesis through careful staging of the process in the developing liver^9,10,23,24^. However, *de novo* lumen formation and expansion occurs within a matter of hours and non-uniformly across the fetal liver, precluding the capture of a high-resolution or quantitative understanding of biliary morphogenesis *in vivo*. Mouse models of FGFR-driven ICC have proven difficult to generate and other ICC models feature tumors that develop after relatively long durations and with unpredictable timing and anatomical distribution, also precluding a high-resolution understanding of how they initiate^25,26^. 3D organoids are widely used to model some aspects of epithelial biology, but most biliary organoids established from normal or tumor tissue are spherical cysts that neither form similarly to or recapitulate biliary tubular architecture^27–30^. To study biliary morphogenesis and the impact of normal and oncogenic

FGFR on that process at high resolution, we adapted a physiological 3D model for quantitative imaging and deployed it to analyze a panel of hepatoblast cell lines that uniquely capture discrete stages of *de novo* lumen formation and expansion. We discovered that FGFR signaling is important for normal biliary morphogenesis and that oncogenic versions of FGFR2 disrupt the process; however, we found that the signaling and cellular phenotypes triggered by oncogenic FGFR2 depend on the epithelial maturation of the cells. Surprisingly, we also found that the morphogenetic consequences of different forms of genomically activated FGFR2 are completely distinct, largely proliferation-independent, and driven by ‘nonclassical’ FGFR2 signaling, which has important therapeutic implications. This work illustrates how morphogenesis can inform our understanding of tumor biology and of how tumor-driving mutants can be important tools for studying morphogenesis.

## Results

### Panel of mouse hepatoblasts captures stages of de novo *lumen competence*

To dissect the process of *de novo* lumen formation and extension, we established a panel of 10 clonal hepatoblast (HB) cell lines from E14.5 mouse embryos and assessed their ability to form and extend lumens in a modified 3D collagen sandwich assay (Fig. 1A; Supplemental Fig. 1A)^31^. Although all express the biliary lineage markers *Hnf1b* and *Sox9* (Supplemental Fig. 1B), we found that our HB lines reproducibly exhibit and maintain one of three levels of lumen competence: HB^A^, lumen-incompetent (N = 4 models); HB^B^, immature lumen-competent - many small lumens (N = 4 models); and HB^C^, mature lumen-competent - large interconnecting lumens and tubular structures of ∼10-15μm diameter that mimic small peripheral biliary ductules (N = 2 models)^32^, representative examples of which are shown in Fig. 1A and Supplemental Fig. 1A. For comparison, we deployed NIH3T3 fibroblasts that do not form lumens, and two mature epithelial cells: human cholangiocytes derived from intrahepatic cholangiocyte organoids (ICO)^33^ that form larger diameter tubes (30-60μm) that model interlobular ducts, and Caco2 colonic epithelial cells that instead form lumen-containing spherical cysts (Supplemental Fig. 1C,D). Note that throughout we will use the term ‘epithelial state’ to refer to differences in architectural maturity (ie polarity, junctional and apical surface elaboration, lumen diameter and interconnection, columnar cellular morphology) rather than differentiation, which implies additional knowledge of epigenetics, cell fate and/or function, although these likely correlate. As in developing livers, *de novo* lumens formed by HB cells in 3D initiate within E-cadherin-positive cell-cell junctions and are marked by a concentrated rim of actin and ERM (Ezrin, Radixin, Moesin) proteins that is devoid of E-cadherin (Fig. 1B)^10^. We developed a quantitative fluorescence imaging workflow in which scanning of the entire 3D culture (∼8 mm^2^ chamber) captures thousands of lumens, enabling the rapid measurement of multiple parameters, including lumen number, size and shape (Fig. 1C). We noted that cell division decreased with lumen competence and that lumen extension was enhanced at the 3D chamber periphery, likely due to mechanical forces. Therefore, for each chamber analyzed, all fields quantified were selected to have an equivalent cell density and were spatially located within the outer peripheral 50% (Supplemental Fig. 1E-G). Together, our HB cell panel and quantitative 3D model uniquely capture discrete stages of this pivotal proliferation-independent morphogenetic process.

**Figure 1.**
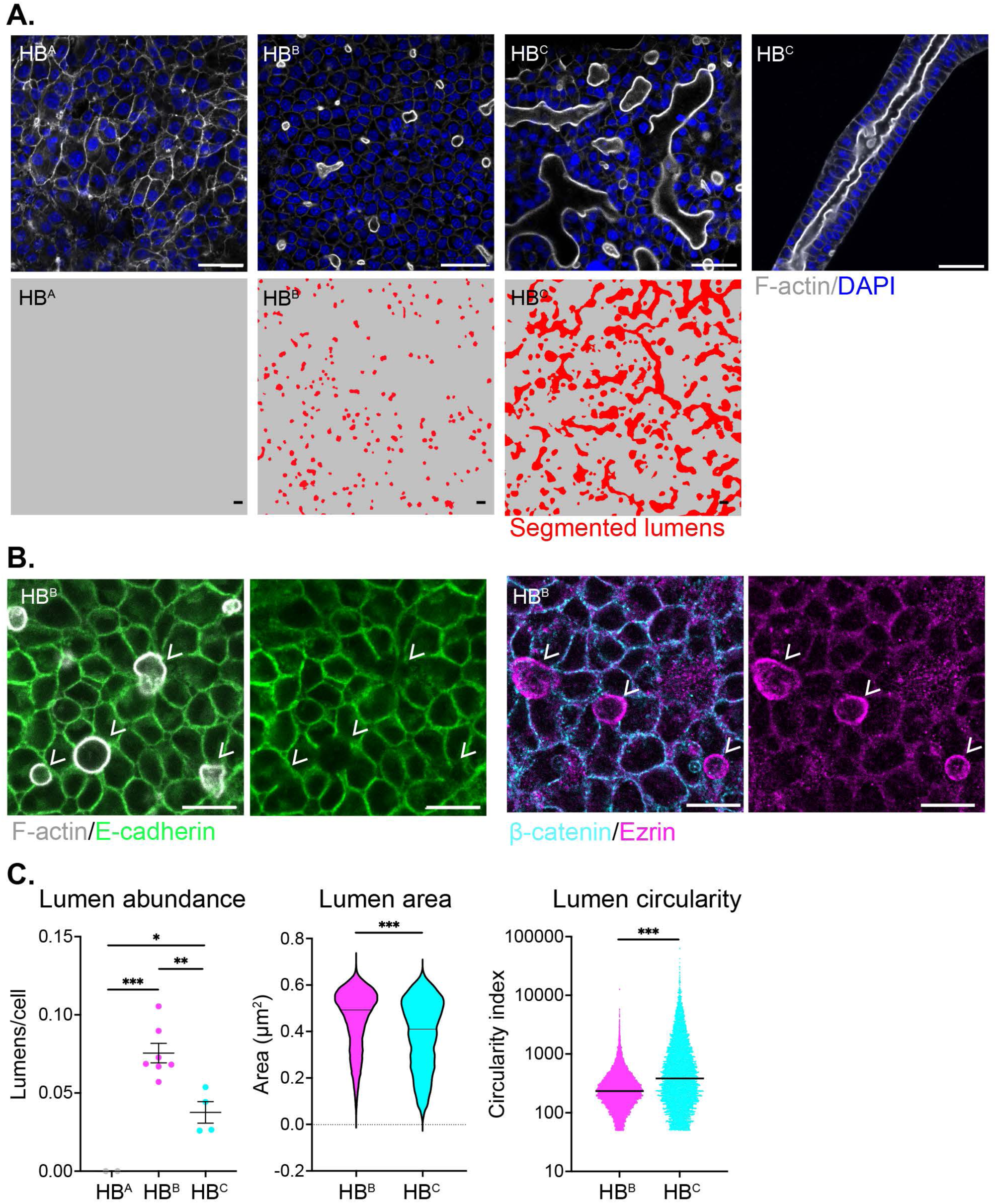
Quantitative imaging of a panel of hepatoblast cell lines that capture discrete stages of lumen competence in a physiological 3D model. A. Confocal images (top) and HALO-based segmentation (bottom) of lumens formed by HB of each class. B. Confocal images of individual lumens (white arrowheads) formed by HB^B^ cells and stained with phalloidin (F-actin, white) and E-cadherin (green) or β-catenin (cyan) and Ezrin (magenta). C. HALO-based quantitation of lumen abundance (lumens/cell; left), size (lumen area; middle) and extension (lumen circularity; right). Bars represent the SEM +/− mean. P values were calculated using a one-way ANOVA with Tukey’s multiple comparisons test (lumen abundance, *p<0.05; ***p<0.001, ns = not significant) or Mann-Whitney test (lumen area and circularity, ***p<0.001). For lumen abundance, each datapoint represents the scan of an entire 3D culture chamber with 0 (HB^A^), 25,981 (HB^B^) or 7,231 (HB^C^) lumens. For lumen area and circularity each datapoint represents an individual lumen. Scale bars = 20 μm.

### Lumen competence coincides with junctional maturation and apical gene expression

Our HB cell lines express similar levels of N-cadherin (*Cdh2*) and E-cadherin (*Cdh1*) protein and mRNA, and no clear transcriptional mesenchymal-to-epithelial transition (MET) accompanies the acquisition of lumen competence (Fig. 2A; Supplemental Fig. 2). However, inspection of cell-cell junctions revealed clear evidence of junctional maturation (Fig. 2B,C). In contrast to the punctate perijunctional E-cadherin and strong linear junctional N-cadherin exhibited by lumen- incompetent HB^A^ cells, immature lumen competent HB^B^ cells form junctions with linear E-cadherin and reduced N-cadherin, while mature lumen-competent HB^C^ cells exhibit condensed linear E-cadherin, a complete loss of junctional N-cadherin, the appearance of the tight junction component ZO-1 without increased *ZO-1/Tjp1* mRNA, and a marked increase in cell height (Fig. 2B-D; Supplemental Fig. 2).

**Figure 2.**
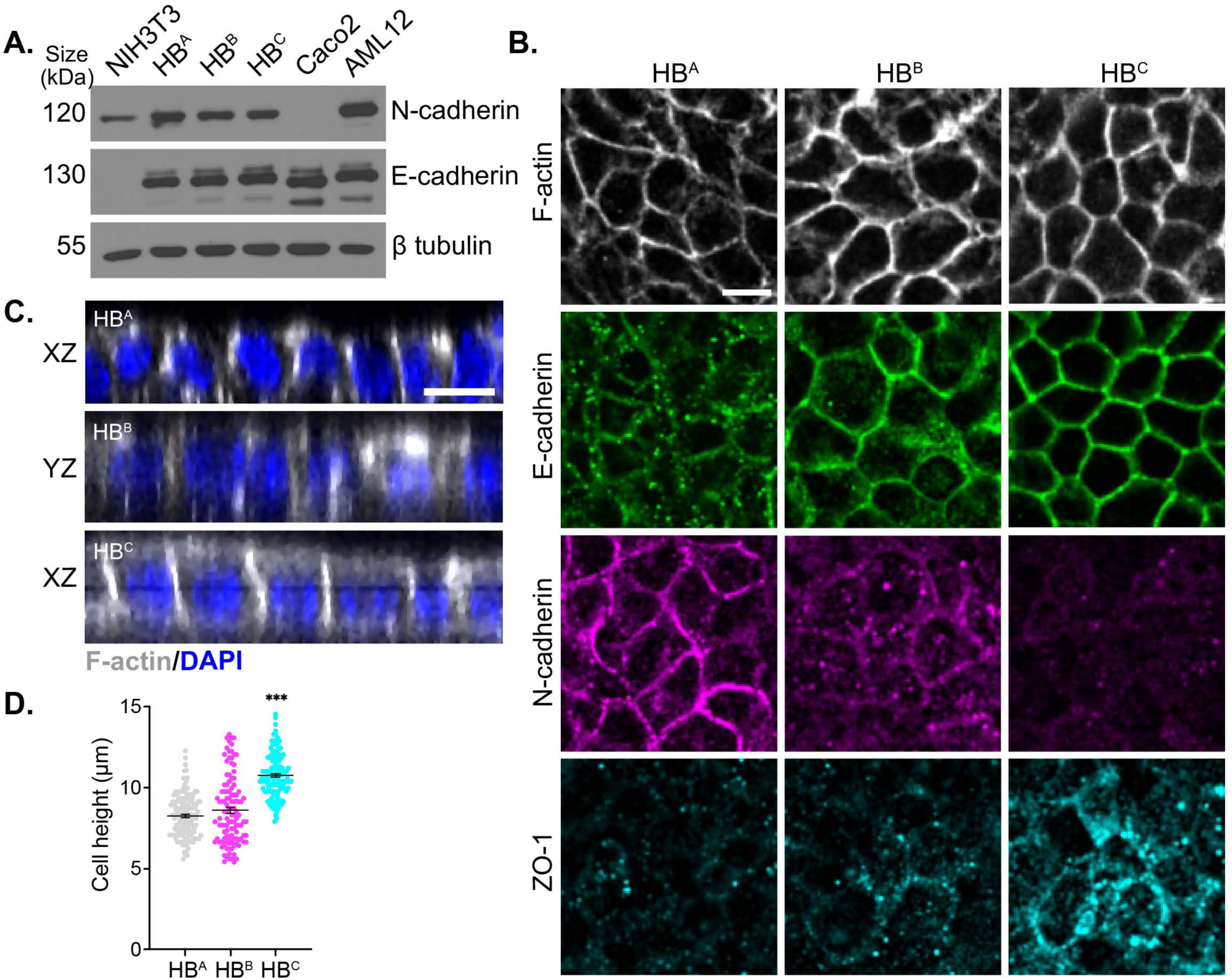
Progressive junction maturation accompanies the ability to form immature and mature lumens. A. Immunoblot showing E-cadherin and N-Cadherin levels across our panel of HBs, NIH3T3, Caco2 and AML12 cells. β-tubulin was used as a loading control. B. Confocal images depicting F-actin (white), E-cadherin (green), N-cadherin (magenta) and ZO-1 (cyan) localization to cell-cell junctions in HB^A^, HB^B^ and HB^C^ cells that were fixed without the addition of the overlay gel. C. Vertical *x-z* and *y-z* views of F-actin depict cell height in HB^A^, HB^B^ and HB^C^ cells. D. Quantitation of cell height from C. P values were calculated using a one-way ANOVA with Tukey’s multiple comparison’s test (***p<0.001). Scale bars = 10 μm.

To understand how HBs acquire lumen competence we compared gene expression profiles across our cell panel by RNAseq (Fig. 3A). We found that lumen-competent HB^B^ and HB^C^ cells, in contrast to lumen-incompetent HB^A^ cells, exhibit a striking increase in the expression of apical membrane components, including transporters and cell junction components. Notably, some apical membrane components exhibit a progressive increase in expression with lumen initiation and maturation, and others stage-specific increases (Fig. 3B-D; Supplemental Fig. 3). For example, genes linked to bile acid metabolism are upregulated in association with lumen initiation (HB^A^→HB^B^) and apical junctional components, including tight junction components, are upregulated in association with lumen maturation, consistent with our observations of junctional maturation (HB^B^→HB^C^ transition; Fig. 3C,D; Fig. 2B). Interestingly, however, principal component analysis (PCA) of the entire panel of HB cell lines revealed that while cell lines of each subtype clustered well, transcriptional differences were much greater between lumen incompetent and competent cell lines, suggesting that posttranscriptional differences play a major role in enabling lumen extension and tubulogenesis (Fig. 3E). Thus, biliary epithelial cell states can be uniquely defined through a combination of gene expression signatures, junctional maturation and lumen-forming capability.

**Figure 3.**
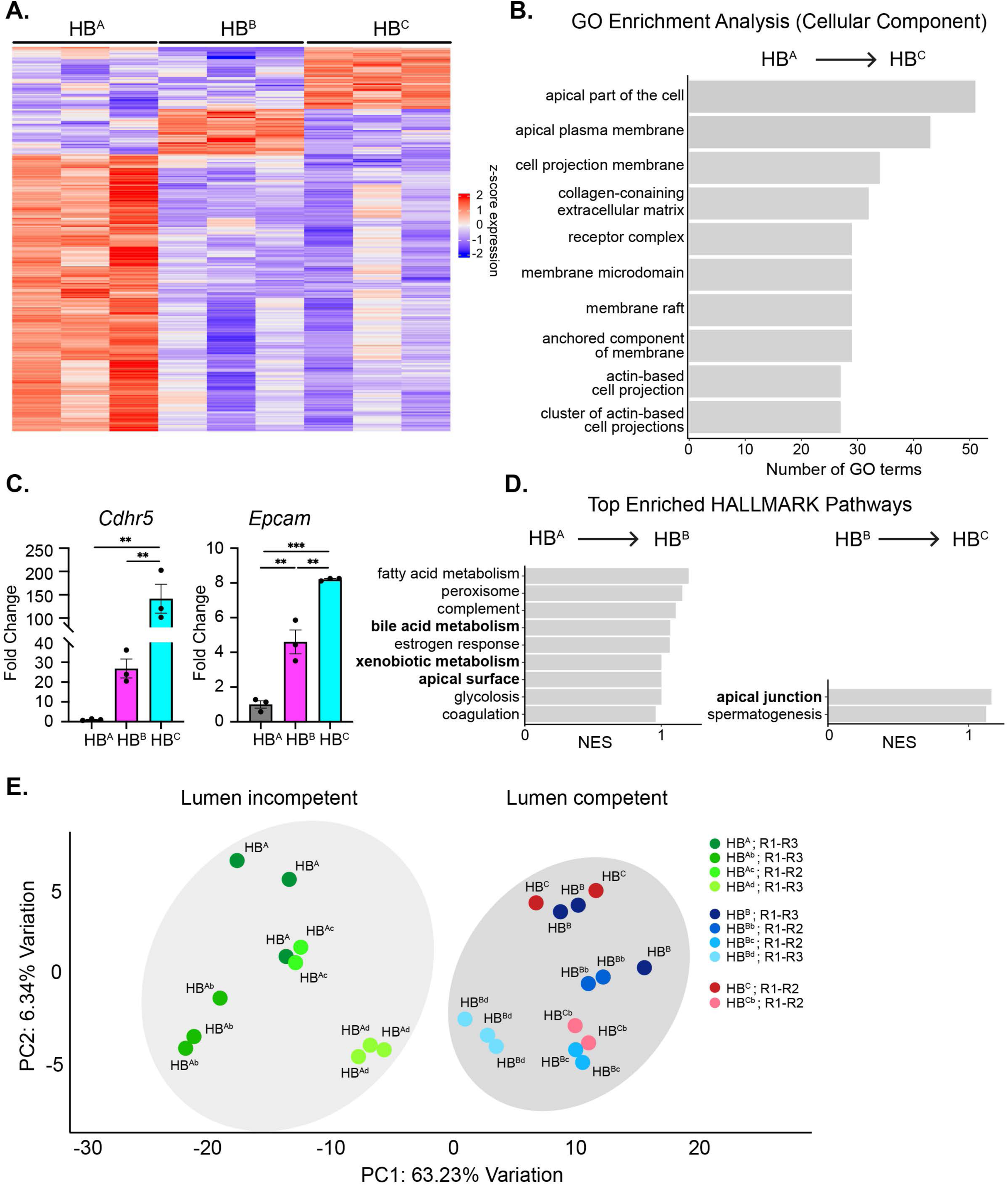
RNAseq reveals that the acquisition of lumen competence is accompanied by increased apical gene expression. A. Heatmap showing hierarchical clustering-based RNAseq profiles of representative HB cell lines of each class of lumen competence. Data is presented as z-scaled log2 normalized expression. Grouped columns reflect RNAseq profiles of three *independent passages* of each cell line as opposed to technical replicates. B. Top Gene Ontology (GO) Cellular Components that are upregulated in mature lumen competent (HB^C^) versus lumen incompetent (HB^A^) HB cells (p < 0.01, q < 0.01). C. Validation of the progressive increase of selected apical gene mRNAs with lumen competence by qPCR. Bars represent the mean +/− SEM. P values were calculated using a one-way ANOVA with multiple comparisons test (*p<0.05; **p<0.01; ***p<0.001). D. Bar plots showing the normalized enrichment scores (NES) for the top Hallmark gene sets that are upregulated in association with lumen initiation (HB^B^ versus HB^A^; left) or lumen extension (HB^C^ versus HB^B^; right). E. Principal component analysis showing clustering of 10 HB cell lines. Each cell line is color coded according to the categorization analysis carried out in Supplemental Figure 1A (HB^A^-, HB^B^-, or HB^C^-like) is and represented by two or three biological replicates, labeled as R1, R2, or R3).

### FGFR signaling promotes de novo lumen formation and biliary morphogenesis

In other examples of *de novo* lumen formation in development, FGFR signaling plays an important role by promoting epithelial character in a manner that is spatially restricted by the *de novo* lumens themselves^13,14^. To determine whether FGFR activity is required for *de novo* lumen formation during biliary morphogenesis, we treated 3D cultures of lumen competent HB^B^ cells with the clinically active FGFR1-3 kinase inhibitor infigratinib (FGFRi). We found that FGFRi treatment of HB^B^ cells significantly decreased lumen number (Fig. 4A,B). Moreover, FGFRi treatment reduced the expression of apical genes associated with the acquisition of lumen competence (Fig. 4C). On the other hand, increasing FGFR signaling via exogenous expression of FGFR2^wt^ led to the upregulation of a set of apical genes that significantly overlaps with those naturally upregulated in lumen-competent HB cells (Fig. 4D,E). These data demonstrate that FGFR signaling promotes *de novo* lumen formation and enhances polarity and epithelial character during biliary morphogenesis as in other developmental settings.

**Figure 4.**
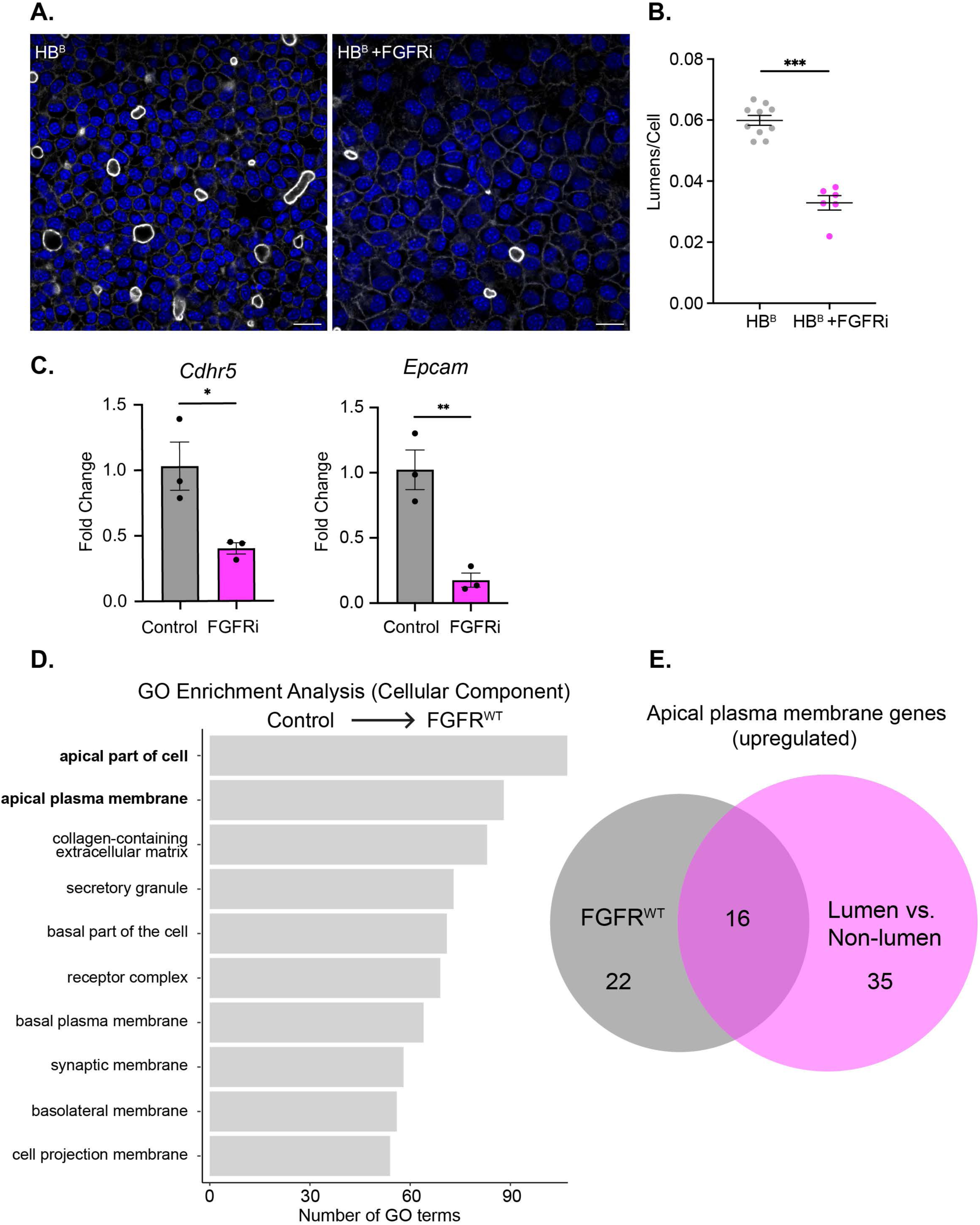
FGFR promotes *de novo* lumen formation and biliary morphogenesis. A. Confocal images of HB^B^ cells in 3D cultures grown without and with the FGFRi infigratinib (3 μM) and stained with phalloidin (F-actin; white) and DAPI (blue). Scale bars = 20 μm. B. HALO- mediated quantitation of lumen abundance in control and FGFRi-treated cultures. C. Measurement of apical gene expression in control and FGFRi-treated HB^B^ cells by qPCR. D. Bar plots depicting the top GO Cellular Components that are upregulated by FGFR2^wt^ expression in HB^B^ cells (p < 0.01, q < 0.01). E. Venn diagram depicting the number of overlapping genes from the “apical plasma membrane” GO term (GO:0016324) whose expression is upregulated by HBs in association with lumen competence (magenta) versus those promoted by FGFR2^wt^ overexpression (gray). Bars represent mean +/− SEM. P values were calculated using an unpaired two-tailed Student’s t-test (*p<0.1,**p<0.01,***p<0.001).

### Distinct localization and phenotypic consequences of ICC-causing FGFR2 mutants

FGFR2 is genomically activated by gene fusions, truncations, extracellular domain point mutations, and extracellular domain in-frame deletions (EIDs) in ICC^34^, each of which is thought to render the receptor ligand-independent (Fig. 5A)^18,35^. We examined the impact of expression of an FGFR2^fusion^ (FGFR2^PHGDH^) or FGFR2^EID^ (FGFR2^WI290>C^) allele on biliary epithelial architecture in HB^B^ cells. While neither variant had a strong effect on proliferation, the two FGFR2 mutants yielded distinct phenotypic consequences in our physiological 3D model (Supplemental Fig. 5A; Fig. 5B-E). The FGFR2^EID^ caused a modest increase in *de novo* lumen number but not size while the FGFR2^fusion^ abolished *de novo* lumen formation (Fig. 5B,C; Supplemental Fig. 5B). Close inspection revealed that the FGFR2^fusion^ but not the FGFR2^EID^ markedly altered E-cadherin cell junctions in which *de novo* lumens form; instead of the discrete linear cell-cell boundary of control and FGFR2^wt^-expressing HB^B^ cells, E-cadherin exhibited a punctate vesicular distribution around cell-cell boundaries in FGFR2^fusion^-expressing cells (Fig. 5D,E). Notably, this effect was specific to E-cadherin, the FGFR2^fusion^ did not impact junctional actin or N-cadherin (Fig. 5B; Supplemental Fig. 5C,D). The biliary marker *Sox9* was not decreased (rather showing a trend toward increased expression), suggesting that the FGFR2^fusion^ did not trigger biliary dedifferentiation (Supplemental Fig. 5E). Importantly, both junctional E-cadherin integrity and lumen formation in FGFR2^fusion^-expressing cells were rescued by FGFRi treatment (Fig. 5B-E). The differential consequences of the two mutants were further underscored by the distinct changes in gene expression they elicited; nearly all of the genes upregulated by the FGFR2^EID^ were also induced by FGFR2^wt^, while the FGFR2^fusion^ induced a largely non-overlapping set of genes (Fig. 5F; Supplemental Fig. 5F).

**Figure 5.**
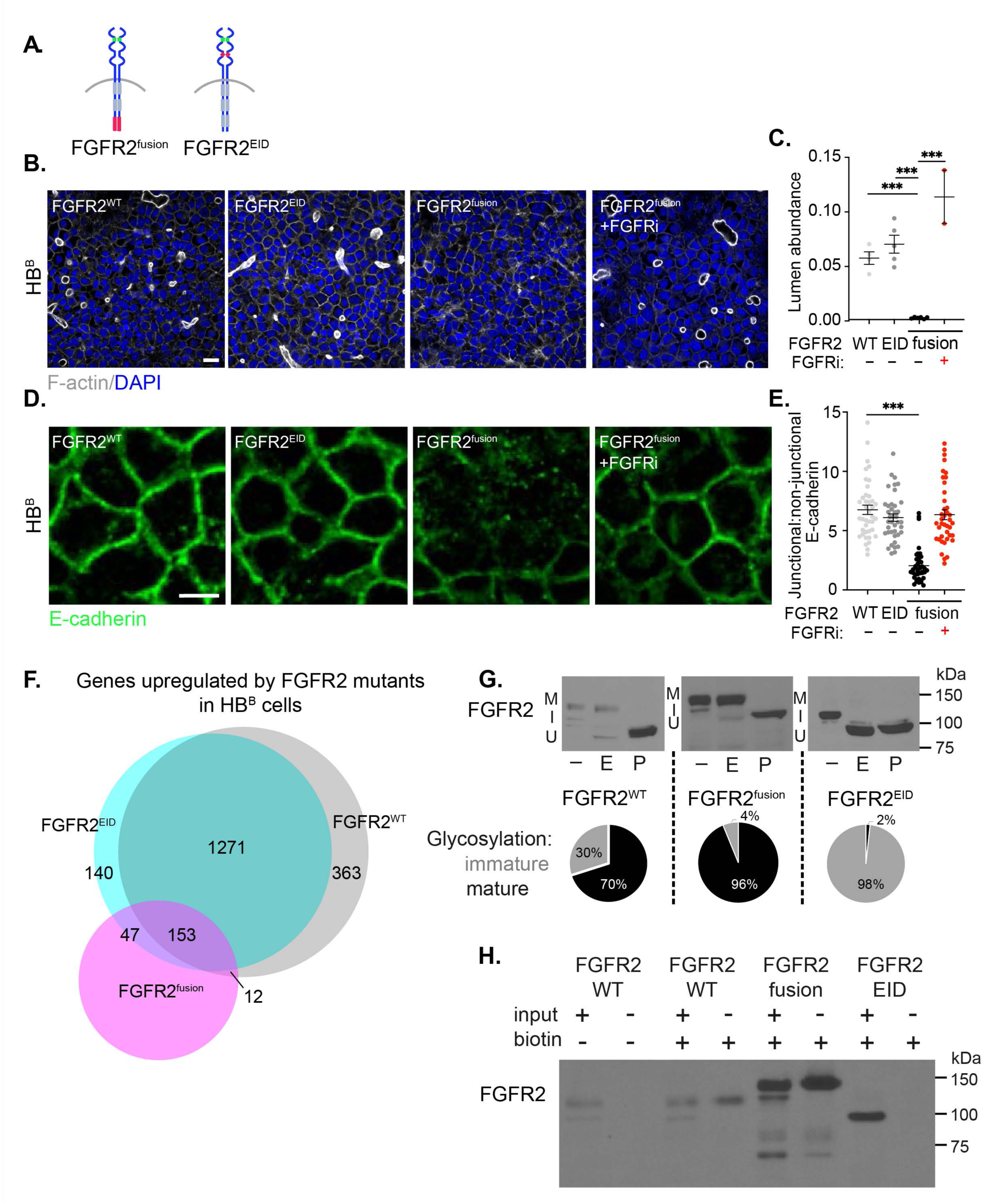
Distinct localizations and phenotypic consequences of FGFR2^fusion^ and FGFR2^EID^ in HB^B^ cells. A. Graphic of the two mutant FGFR2 isoforms with the locations of the fused segment and extracellular domain mutations in red. B. Confocal images depicting lumen formation (F-actin, white) in HB^B^ cells expressing FGFR2^wt^, FGFR2^EID^, FGFR2^fusion^ or FGFR2^fusion^ plus FGFRi. C. Quantitation of lumen abundance across the entire 3D chambers from (B) plus at least one biological replicate. D. Confocal images depicting E-cadherin based junctions (green) in HB^B^ cells expressing FGFR2^wt^, FGFR2^EID^, FGFR2^fusion^ or FGFR2^fusion^ plus FGFRi. E. Quantitation of junctional integrity in 3D cultures of HB^B^s expressing FGFR2^wt^, FGFR2^EID^ and FGFR2^fusion^ with and without FGFRi. F. Venn diagram depicting overlapping genes upregulated by FGFR2^wt^, FGFR2^EID^ and FGFR2^fusion^ in HB^B^ cells. G. *Top*, Immunoblot of lysates from FGFR2^wt^-, FGFR2^EID^- and FGFR2^fusion^-expressing HB^B^ cells treated with EndoH (E) or PNGase (P). M = mature, I = immature, U = unglycosylated. *Bottom*, Pie chart representation of the proportion of immature and mature glycosylated forms of each version of FGFR2. H. Immunoblot of total and surface biotinylated proteins from FGFR2^wt^, FGFR2^fusion^ and FGFR2^EID^- expressing cells. Bars represent mean +/− SEM. P values were calculated using a one-way ANOVA with Tukey’s multiple comparisons test (***p<0.001).

We next compared the processing and subcellular localization of the FGFR2 variants. Immunoblotting revealed that in HB^B^ cells the majority of FGFR2^wt^ is fully glycosylated and sensitive to peptide-N-glycosidase F (PNGase F), which removes all asparagine-proximal N-linked glycans, whereas ∼30% remains in an immature state of glycosylation that is uniquely sensitive to treatment with Endoglycosidase H (EndoH) (Fig. 5G). On the other hand, the FGFR2^fusion^ is fully glycosylated with little detectable EndoH-sensitive receptor population (Fig. 5G). In contrast, nearly all of the FGFR2^EID^ (∼98%) is in an immature state of glycosylation (Fig. 5G). Immunofluorescence revealed that the FGFR2^fusion^ is concentrated at cell-cell boundaries, while the FGFR2^EID^ is largely excluded and is instead trapped within the ER-Golgi in HB^B^ cells, as has been described in NIH3T3 cells (Supplemental Fig. 5G)^35^. Surface biotinylation confirmed the enrichment of FGFR2^fusion^ at the surface, while surface FGFR2^EID^ was undetectable (Fig. 5H). Thus, in immature lumen-forming HB^B^ cells, two ICC-causing FGFR2 variants are trafficked differently and elicit distinct phenotypic consequences.

### Glycosylation and surface availability of FGFR2^EID^ depend on epithelial state

Our HB cells exhibit a range of epithelial character, so we asked whether FGFR2^EID^ glycosylation varied across this biliary-relevant panel. We found that, as in NIH3T3 and HB^B^ cells, the FGFR2^EID^ is immaturely glycosylated and ER-Golgi-trapped in lumen-incompetent HB^A^ cells (Fig. 6A). However, in mature lumen-competent HB^C^ cells, a significant proportion of the

**Figure 6.**
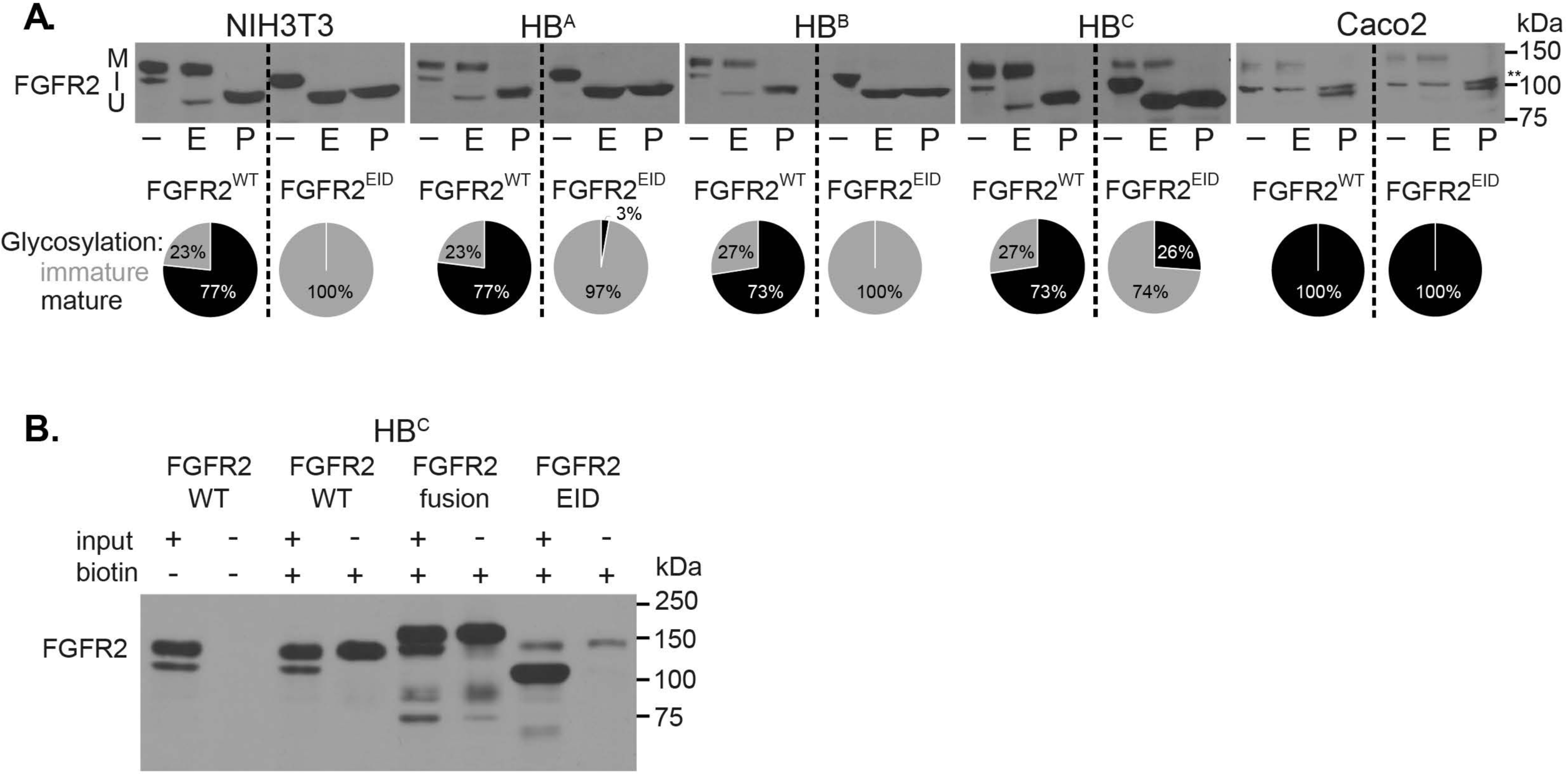
Glycosylation and trafficking of FGFR2^EID^ depends on epithelial character. A. *Top*, immunoblots of lysates from NIH3T3 fibroblasts, HB^A^, HB^B^ and HB^C^ HBs and Caco2 epithelial cells treated with EndoH (E) or PNGase (P). M = mature, I = immature, U = unglycosylated. * = irrelevant human-specific background band. *Bottom*, Pie charts representing the proportions of immature (gray) and mature (black) glycosylated forms of each version of FGFR2. B. Immunoblot showing input and surface biotinylated FGFR2^wt^, FGFR2^fusion^ and FGFR2^EID^ levels in HB^C^ cells.

FGFR2^EID^ is glycosylated (26%) and reaches the surface while in even more mature Caco2 epithelial cells the FGFR2^EID^ is fully glycosylated (Fig. 6A,B). Notably, the FGFR2^EID^ also exhibited immature glycosylation in murine AML12 hepatocytes, suggesting that mature glycosylation of this mutant is specific to monopolar columnar epithelial cells (Supplemental Fig. 6A). Finally, we found that a second FGFR2^EID^ (FGFR2^H167_N173^) was also entirely immaturely glycosylated in NIH3T3 and HB^B^ cells but maturely glycosylated in Caco2 cells (Supplemental Fig. 6B)^18^. Therefore, the trafficking of ICC-causing FGFR2^EID^ mutants depends on biliary epithelial state.

### Epithelial state-specific consequences of FGFR2^EID^ and FGFR2^fusion^

As both FGFR2^EID^ and FGFR2^fusion^ can be maturely glycosylated and reach the surface in HB^C^ cells that form mature junctions and extended lumens (Fig. 6), we asked whether they cause similar phenotypes in these cells. We found that the two mutants again elicited distinct changes in gene expression (Fig. 7A). Moreover, while FGFR2^EID^-induced gene expression changes mirror those of FGFR2^wt^ in HB^C^ cells, as they did in HB^B^ cells (Fig. 7A, 5A), the programs triggered by FGFR2^EID^/FGFR2^wt^ are epithelial state-dependent, while those triggered by FGFR2^fusion^ are similar in HB^B^ and HB^C^ cells (Supplemental Fig. 7A). Notably, FGFR2^EID^/FGFR2^wt^ again upregulate apical gene expression in HB^C^ cells, while the FGFR2^fusion^ instead *downregulates* a largely non-overlapping set of apical genes (Supplemental Fig. 7B,C). The FGFR2^EID^ and FGFR2^fusion^ also elicited distinct and epithelial state-specific changes in 3D morphology. As is evident in low magnification images the phenotypic consequences of both mutants depend on the epithelial maturity of the cell. For example, in contrast to the specific disruption of junctional E-cadherin in HB^B^ cells, expression of the FGFR2^fusion^ in HB^C^ cells caused striking aggregation of both E-cadherin and actin near multicellular junctions but had no discernable effect in HB^A^ cells that exhibit punctate E-cadherin distribution and preferentially form N-cadherin based junctions at baseline (Fig. 2B; Supplemental Fig. 7D,E). These junction phenotypes were accompanied by distinct impacts on *de novo* lumen formation and extension; instead of completely blocking *de novo* lumen formation as in HB^B^ cells, the FGFR2^fusion^ converted the large extended and interconnected lumens of HB^C^ cells to many small, poorly organized lumens (Fig. 1A,C; Fig. 5 B,C; Fig. 7D). Examination of the *z*-plane revealed that while lumens in control HB^C^ cells were centered and aligned vertically, lumens in FGFR2^fusion^- expressing HB^C^ cells were randomly oriented and often remained intracellular, suggesting a lack of spatially coordinated polarity (Fig. 7C,D). In mature epithelial Caco2 cells FGFR2^fusion^ also impeded coordinated polarity, yielding cysts containing multiple lumens (Supplemental Fig. 7F). Thus, despite enhanced surface availability in all HB cells, the phenotypic consequences of FGFR2^fusion^ depend on epithelial maturity.

**Figure 7.**
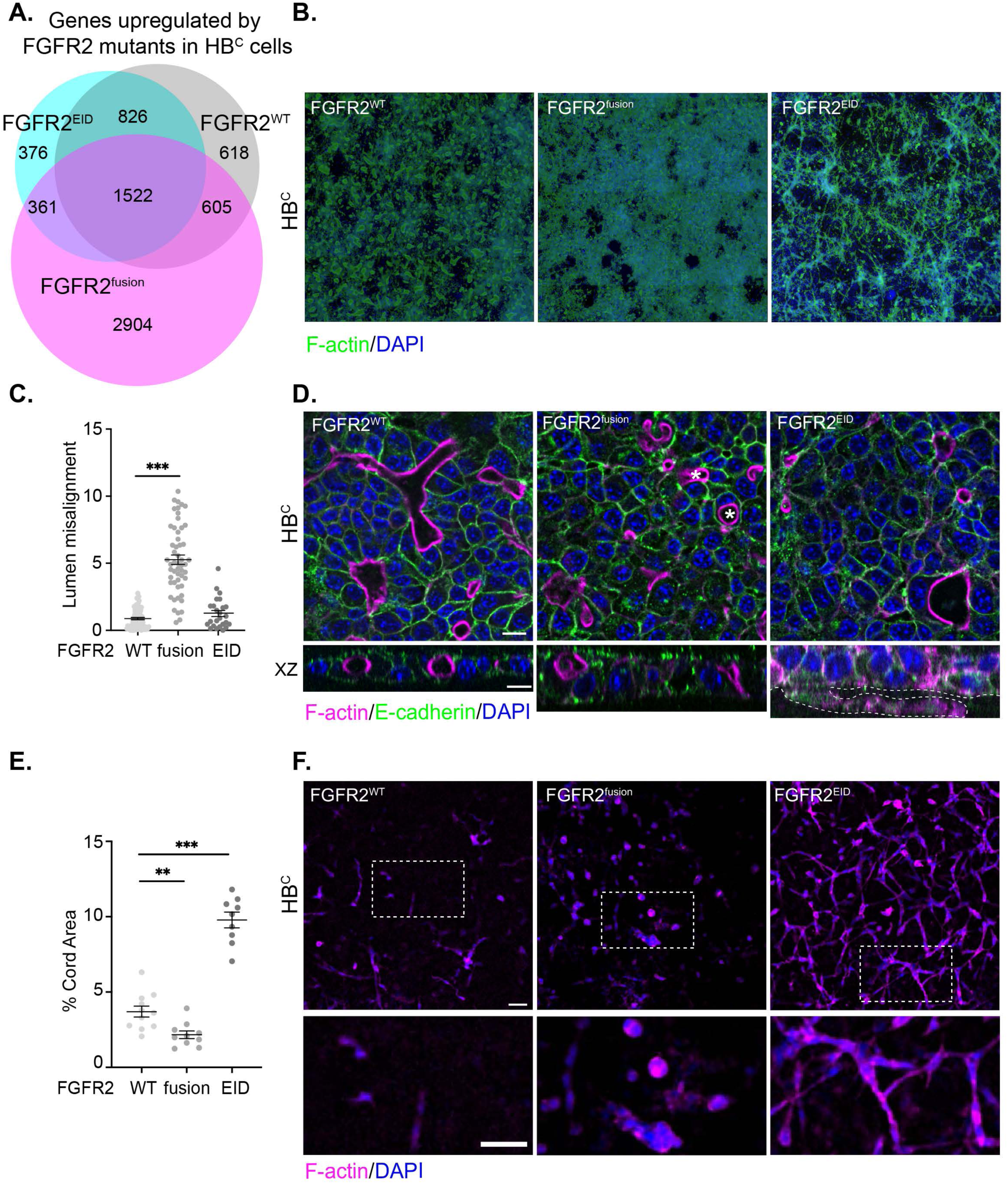
Differential impact of FGFR2^EID^ and FGFR2^fusion^ on lumen formation and morphogenesis in lumen-extending HB^C^ cells. A. Venn diagram showing overlapping numbers of genes upregulated by FGFR2^wt^, FGFR2^EID^ or FGFR2^fusion^ in HB^C^ cells. B. Low power immunofluorescence images of the entire 3D culture of FGFR2^wt^-, FGFR2^fusion^- or FGFR2^EID^- expressing HB^C^ cells stained for F-actin (green) and DAPI (blue). C. Quantitation of the area of cords in FGFR2^wt^-, FGFR2^fusion^-, or FGFR2^EID^-expressing HB^C^ cells. D. Confocal images of a z- plane basal to the cell monolayer in FGFR2^wt^-, FGFR2^fusion^- or FGFR2^EID^-expressing HB^C^ cells stained for F-actin (magenta) and DAPI (blue). Scale bars = 50 μm. E. Quantitation of lumen alignment in FGFR2^wt^-, FGFR2^fusion^- and FGFR2^EID^-expressing HB^C^ cells. F. *Top*, x-y and *bottom*, x-z confocal images of F-actin (magenta), E-cadherin (green) and DAPI (blue) stained 3D cultures of FGFR2^wt^-, FGFR2^fusion^- and FGFR2^EID^-expressing HB^C^ cells. Dashed line depicts the basal cord emerging from the cell monolayer in FGFR2^EID^-expressing HB^C^ cells. Scale bars = 10 μm. Bars represents mean +/− SEM. P values were calculated using a one-way ANOVA with Tukey’s multiple comparisons test (**p<0.01, ***p<0.001).

In contrast to the FGFR2^fusion^, the FGFR2^EID^ did not appreciably affect E-cadherin junctions in any of our cells but impacted lumen formation in a way that was both distinct from the FGFR2^fusion^ and dependent on epithelial maturity. In contrast to HB^B^ cells, where the FGFR2^EID^ promoted a modest increase in lumen formation, the FGFR2^EID^ impeded lumen extension in HB^C^ cells without altering their spatial distribution (Fig. 5B,C; 7B-D). In Caco2 cells the FGFR2^EID^ had no discernable impact on cyst formation (Supplemental Fig. 7F).

Notably, the two tumor-causing FGFR2 mutants also triggered distinct forms of ECM invasion from the basal surface of the lumenized HB^C^-formed biliary structures. FGFR2^fusion^-expressing cells formed small spherical cell clusters that emerged from the basal surface and exhibited an actin-rich outer surface, consistent with an amoeboid mode of invasion (Fig. 7F)^36^. Conversely, FGFR2^EID^-expressing cells formed long cords of cells extending from the basal surface, explaining their strikingly different appearance at low magnification (Fig. 7B,E,F). Altogether these data indicate that two ICC-driving FGFR2 mutants differentially impact both key aspects of biliary architecture and invasive tendency in a manner that depends on biliary epithelial state and could have profound impacts on how tumors initiate and progress.

### Proliferation- and MAPK-independent phenotypic consequences of the FGFR2^fusion^ and FGFR2^EID^ in mature epithelial cells

Activation of FGFR triggers signaling through multiple pathways, but phosphorylation of the adaptor scaffold FGF receptor substrate 2 (FRS2) and consequent MAPK pathway activation is thought to be primarily responsible for the oncogenic consequences of mutant FGFR^18,37^.

Despite having profound effects on 3D architecture, neither FGFR2 variant significantly increased the proliferation of HB^B^ or HB^C^ cells (Supplemental Fig. 5A, 8A). In HB^B^ cells, the FGFR2^fusion^ triggered strong tyrosine phosphorylation of FRS2 and activation of the MAPK pathway effector MEK, but the FGFR2^EID^ did not activate any of them (Fig. 8A). This was surprising given that the same FGFR2^EID^ has been shown to activate FRS2-MAPK signaling and transform NIH3T3 cells despite being immaturely glycosylated and ER-Golgi-trapped, which we verified (Fig. 6A, Supplemental Figure 8B)^18,35^. However, the FGFR2^EID^ also did not activate FRS2 in mature lumen-competent HB^C^ cells where it was partially glycosylated and had strong morphogenic consequences (Fig. 8A, 6A,B, 7). Also surprisingly, in contrast to NIH3T3, HB^A^ and HB^B^ cells, the FGFR2^fusion^ activated FRS2-MAPK only modestly in HB^C^ cells despite profoundly impacting their architecture (Fig. 8A, 7). These observations suggest that signaling other than MAPK plays an important role in the three-dimensional phenotypes elicited by the two mutants. Indeed, while FGFR inhibition completely rescued FGFR2^fusion^- and FGFR2^EID^-elicited 3D phenotypes, MAPK kinase inhibition (MEKi) did not (Fig. 8B-D). Notably, immunoblotting revealed a reduction in size and increase in tyrosine phosphorylation of FRS2 in MEKi treated FGFR2^fusion^- and FGFR2^EID^-expressing HB^C^ cells, consistent with the reported relief of negative feedback seen in other cell types (Fig. 8E)^38^. While MEKi did not block basal cord formation by FGFR2^EID^-expressing HB^C^ cells, it induced the formation of small intercellular lumens within the cords that were not present in vehicle-treated FGFR2^EID^-expressing cells (Fig. 8D). Instead, pharmacologic inhibition of Src family tyrosine kinases (SFKs), which are key FGFR effectors in multiple developmental contexts and have well-known roles in signaling at the basal surface of epithelial cells, completely rescued the basal cords elicited by FGFR2^EID^ in HB^C^ cells (Fig. 8F, Supplemental Fig. 8C)^39,40^. These observations are consistent with the possibility that FGFR2- FRS2 and FGFR2-SFK signaling can promote apical and basal surface generation, respectively, and that the FGFR2^fusion^ and FGFR2^EID^ differentially hijack these two activities, which could have profoundly different impacts on tumor progression and therapeutic response.

**Figure 8.**
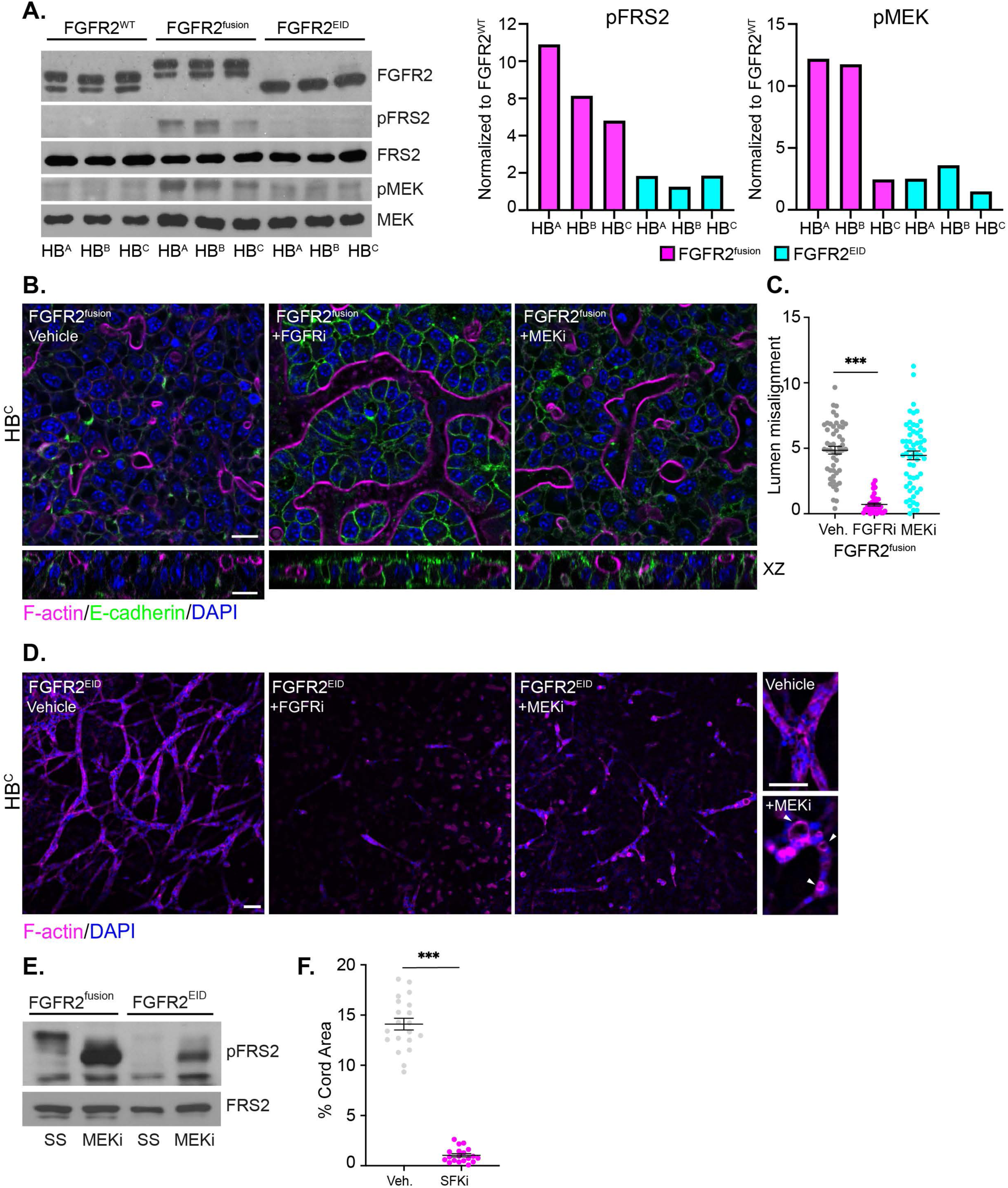
MAPK-independent signaling contributes to FGFR2 mutant 3D phenotypes in mature epithelial cells. A. *Left*, Immunoblots showing FGFR2^wt^-, FGFR2^fusion^- and FGFR2^EID^- induced signaling in HB^A^, HB^B^ and HB^C^ cells. *Right*, Quantitation of the immunoblot bands representing pFRS2 and pMEK normalized to FGFR2^wt^. B. *Top,* x-y, and *bottom,* x-z confocal images of F-actin (magenta), E-cadherin (green) and DAPI (blue) stained 3D cultures of HB^C^ cells expressing FGFR2^fusion^ treated with vehicle (DMSO), FGFRi (3 µM) or MEKi (1µM). C. Quantitation of lumen abundance and alignment in FGFRi- and MEKi-treated FGFR2^fusion^- expressing HB^C^ cells. D. *Left,* Confocal images taken at z planes below the cell monolayer of FGFR2^EID^-expressing HB^C^ cells treated with vehicle (DMSO), FGFRi (3 µM), or MEKi (1 µM). *Right*, detailed views of individual cell cords in vehicle- or MEKi-treated FGFR2^EID^-expressing HB^C^ cells. E. Immunoblot showing pSFK levels in FGFR2^fusion^ and FGFR2^EID^-expressing HB^C^ cells with and without SFKi treatment (1 µM). SS = steady state. F. Quantitation of cord area in vehicle (DMSO) and SFKi-treated (1 µM) FGFR2-EID-expressing HBC cells. Bars represents mean +/− SEM. P values were calculated using a one-way ANOVA with Tukey’s multiple comparisons test (***p<0.001). Scale bars = 10 μm.

### Comparison to human ICC cells

Heterogeneity is a major impediment to developing effective treatments for ICC. Notably, ICC has been classified anatomically into small (peripheral) and large (interlobular) ductal subtypes, but neither the molecular basis or therapeutic relevance of this classification is clear^41,42^. Recent multi-omics analyses also parse human liver cancer cell lines into four major subgroups (R1- R4), with most ICCs belonging to two subgroups – designated ‘bilineage’ (R3) and ‘ductal’ (R4) to reflect dual hepatocyte/bile duct or bile duct-restricted gene signatures, respectively^43^. These signatures stratify primary human ICCs, correlating with clinical outcomes^43^. All FGFR2^fusion+^ ICC human cell lines belong to the R3 subgroup; to our knowledge, FGFR2^EID+^ ICC cell lines have not been established. Transcriptionally, all of our HB lines more closely resemble the R3 rather than R4 ICC subgroups, consistent with the idea that R3 tumors map to a less mature biliary epithelial cell (Fig. 9A). Interestingly the HB lines also express R1 gene signatures, raising the possibility that R1 and R3 tumors have a shared cell of origin. We examined 3 FGFR2^fusion+^ R3 subgroup ICC cell lines, all of which, like our HBs, express both E-cadherin and N-cadherin (Supplemental Fig. 9A). In 3D two of the lines (ICC13-7, ICC21) formed a modest number of small lumens, and for both, FGFRi treatment significantly improved E-cadherin junction integrity and both the number and size of lumens (Fig. 9B-F). Notably, the size of lumens formed by these R3 ICC cells is comparable to those formed by HB cells rather than ICO cells, and to that of peripheral biliary ductules rather than intralobular ducts, suggesting that R3 and R4 subgroups may map to the two anatomical subtypes^43^. In contrast, the third R3 ICC cell line (ICC10-6) did not form lumens with or without FGFRi treatment, despite well-formed E-cadherin junctions (Fig. 9B-F). Importantly, ICC13-7 and ICC21 are known to be sensitive to FGFRi in long-term 2D viability assays while lumen-incompetent ICC10-6 cells are resistant due to high baseline EGFR activity^37,43^. These data identify lumen-forming capability as a novel biomarker of epithelial state for ICC tumor cells that, for FGFR2^fusion+^ ICCs, may correlate with therapeutic response.

**Figure 9.**
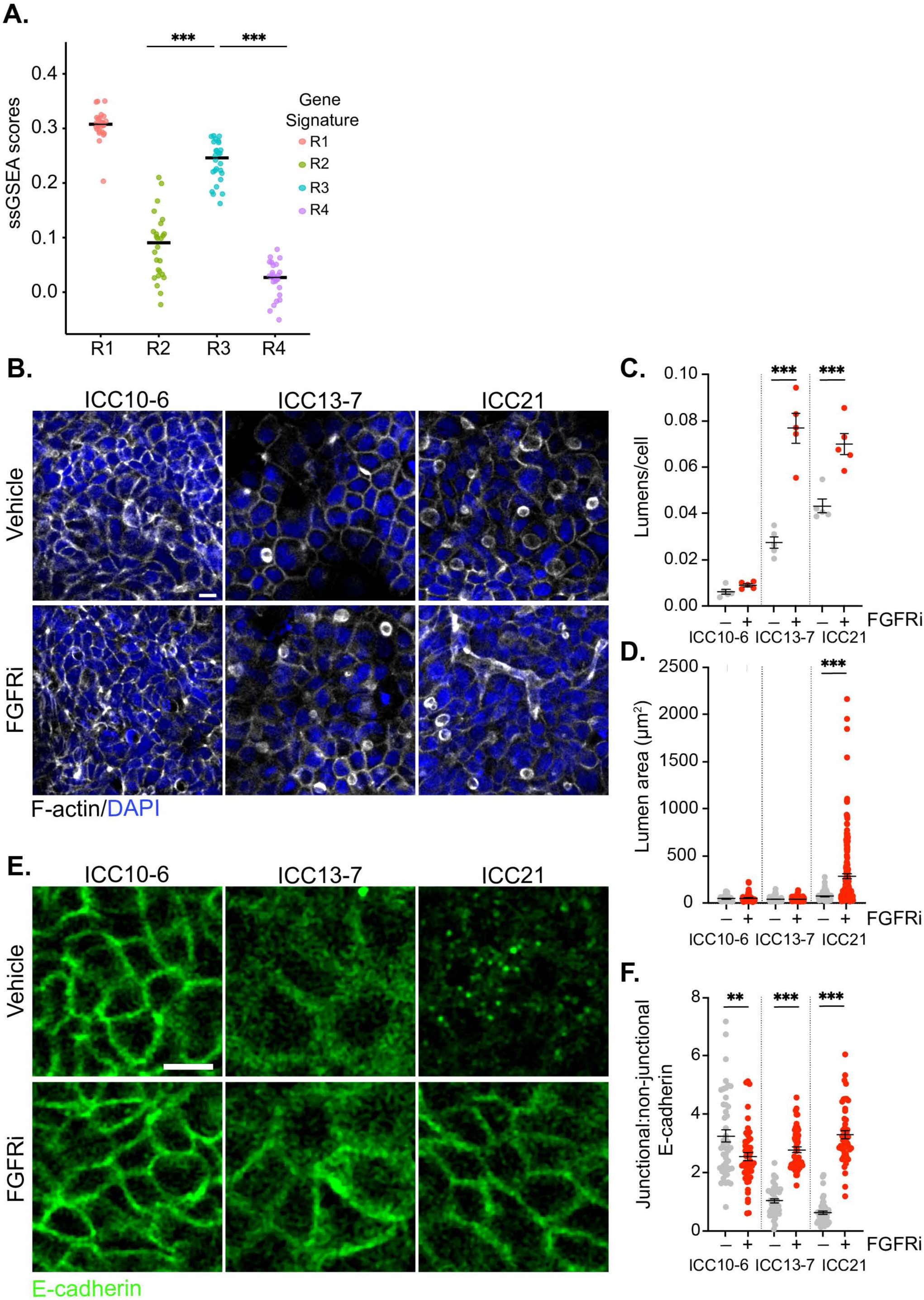
FGFR-dependent architectural defects exhibited by FGFR2^fusion+^ human ICC cells. A. Plot showing single sample GSEA scores for human ICC transcriptional signatures (R1-R4) in HBs. P values were calculated using the Mann Whitney U test (***p<0.001). B. Confocal images depicting lumen formation (F-actin, white; DAPI, blue) in 3D cultures of three FGFR2^fusion+^ human ICC tumor cell lines with and without FGFRi (3 µM). C,D. Quantification of lumen number and area in B. E. Confocal images depicting E-cadherin^+^ junctions (green) in 3D cultures of FGFR2^fusion+^ human ICC cells with and without FGFRi (3 µM). F. Quantitation of junctional E-cadherin in E. P values were calculated using a one-way ANOVA with Tukey’s multiple comparisons test (**p<0.01, ***p<0.001). Scale bars = 10 μm.

## Discussion

Despite major advances in both tissue engineering and tumor bioinformatics, few studies focus on the interface between morphogenesis and tumorigenesis. Our work underscores the facts that tumor-causing mutations often hijack features of morphogenesis in informative ways, that physiologically relevant 3D models of morphogenesis can inform our understanding of tumor biology, and that the morphogenetic capability of tumor cells is an important functional biomarker to contribute to any multi-omics analysis. Organogenesis-based models enable genetic and pharmacologic dissection of normal morphogenetic processes and more accurately model the earliest stages of tumor initiation and progression, the consequences of individual tumor-driving mutations, and drug treatment response. Indeed, our studies have contributed substantially to our understanding of both biliary morphogenesis and tumorigenesis and the role of FGFR activity in both.

By establishing a panel of hepatoblast cell lines and deploying them in a quantitatively adapted physiological 3D model, we uncovered key steps in biliary morphogenesis that would not be captured by single-cell sequencing or other methods. For example, the discovery that lumen initiation and extension are separable events will allow us to study them individually.

Understanding how lumens extend and interconnect to form a functional biliary tubular network is particularly important and timely given the intense effort and recent success in establishing rudimentary hepatobiliary junctions in 3D for liver tissue engineering^44,45^. Beyond physically maturing the apical surface by adding junctional components, cortical scaffolding, receptors and transporters (Fig. 2,3; Supplemental Fig. 3), lumen extension and interconnection requires the dynamic and spatiotemporally appropriate remodeling of apical and junctional surface to enable coordinated alignment of new luminal surfaces among cells. Successful lumen extension and interconnection likely depends on the number and distribution of initiated lumens, which occurs randomly across our 3D cultures (Fig. 1A; Supplemental Fig. 1B)^46^. In contrast to multipolar hepatocytes, developing biliary epithelial cells are strictly monopolar from the onset of polarization. Therefore, when one cell initiates a lumen in partnership with a neighboring cell, neither can participate in lumen formation with a different neighbor. If too many lumens are initiated, rosettes form without lumen extension; this may occur in MEKi-treated HB^C^ cells due to the release of feedback inhibition to FGFR-FRS2, which increases lumen initiation even within FGFR2^EID^-induced basal cords (Fig. 8B,D). On the other hand, initiated lumens that extend and impose lumen formation on uninitiated neighbors can align and connect with other extending lumens. Apical constriction is important for limiting lumen extension in the developing liver and must occur perpendicular to the extending axis, likely due to planar polarized cues^10^. Lumen regression, which is thought to occur *in vivo*, also likely facilitates lumen network formation^10^. A deeper understanding of this dynamic process will be enabled by live-imaging in our 3D model.

The discovery that FGFR signaling is important for biliary *de novo* lumen formation is consistent with its known role in this process in other developmental contexts, and with the reported general requirement for FGFR in biliary morphogenesis in the chick^13,14,47^. Our studies of two different ICC-causing mutants yielded surprising new insights into the trafficking, signaling and cellular functions of FGFR. For example, we found that the FGFR2^EID^ is not fully glycosylated and can signal to FRS2-MAPK from the ER-Golgi in NIH3T3 cells, as has been reported^35^, but not in any of our HB cells or in AML-12 hepatocytes (Fig. 8A), suggesting that liver cells have mechanisms of silencing ER-Golgi-localized FGFR2. Identifying this mechanism will be important given that a significant proportion of FGFR2^wt^ is also in the ER-Golgi in liver and other cell types and relief of this mechanism could be exploited by tumor cells (Supplemental Fig. 5G, Fig. 6A). Our studies also reveal that glycosylation of the FGFR2^EID^ and trafficking to the cell surface *does* occur in more mature epithelial cells (Fig. 6A-B). Thus studies of FGFR2^EID^ uncover epithelial-specific regulation of both FGFR2 biosynthetic transport and membrane signaling, which could explain the lack of epithelial phenotypes or cancer predisposition in congenital craniosynostosis syndromes caused by inherited FGFR2 extracellular domain mutations, including Antley-Bixler and Pfeiffer syndromes, which can be caused by an FGFR2^EID^ mutation nearly identical to that used here (FGFR2^W290C^)^8,48^.

Analysis of the FGFR2^fusion^ also uncovered key functional relationships between FGFR and epithelial biology. The observation that the FGFR2^fusion^ specifically disrupts junctional E-cadherin (but not N-cadherin) by triggering a punctate peri-junctional appearance of E-cadherin is consistent with a role for FGFR in causing E-cadherin endocytosis^49^ and could be an important mechanism of locally removing E-cadherin from cell-cell junctions to initiate and extend lumens. The enhanced surface availability of the FGFR2^fusion^ and ectopic, spatially uncoordinated lumens formed by FGFR2^fusion^-expressing HB^C^ cells suggests that the constitutive dimerization provided by the fusion partner and/or removal of the extreme FGFR2 C-terminal domain compromises the normal spatial restriction of FGFR2 during *de novo* lumen formation (Fig. 5A). Notably, like the FGFR2^EID^, signaling from the FGFR2^fusion^ to FRS2-MAPK is reduced in mature epithelial cells, again suggesting that epithelial-specific mechanisms of dampening canonical FGFR2 signaling exist.

Our studies have several important clinical implications. First, our discovery that FGFR2^fusion^ and FGFR2^EID^ behaved differently by every criterion that we analyzed suggests that they elicit distinct tumor behaviors from initiation to metastasis and may therefore exhibit distinct cooperating genetic events, therapeutic vulnerabilities and resistance mechanisms, microenvironments, and disease progression. This could explain reports that FGFR2^EID+^ and FGFR2^fusion+^ human ICCs differentially respond to FGFRi^50–52^. It will be important to evaluate other types of FGFR2 alteration that occur in ICC and determine whether different FGFR alterations also yield distinct phenotypes in other cancers. Second, although MAPK signaling is widely thought to be the tumor-driving consequence of FGFR mutation in ICC and other cancers, the proximal phenotypic consequences of both the FGFR2^fusion^ and FGFR2^EID^ in our models are proliferation- and MAPK-independent. In fact, the striking release of negative feedback to FGFR2-FRS2 upon MEKi treatment could enhance MAPK-*independent* morphogenetic consequences with important clinical implications given that MEKi are being actively considered for ICC in certain situations^53^. The distinct FRS-MAPK-independent consequences of each mutant underscore the importance of non-canonical FGFR signaling and suggest that tumor-initiating morphogenetic defects may enable aberrant microenvironmental signals and/or cooperating genetic events that synergize to yield MAPK activation and transformation. Recent genetic studies in mice highlight Src as an important MAPK-independent FGFR effector in development and we found that SFKi blocks the basal cords triggered by FGFR2^EID^; it is therefore notable that some ICCs were found to be sensitive to Src inhibition in a recent high-throughput *in vitro* screen^39,54^. Finally, our studies provide early support for lumen-forming capacity as an important and potentially therapeutically predictive biomarker for ICC tumor cell lines. Further studies that exploit the genetic and pharmacologic manipulability of our quantitative physiological 3D model will allow us to fully dissect the distinct mechanisms by which FGFR2^fusion^, FGFR2^EID^ and other ICC-causing alterations influence morphogenesis and tumorigenesis.

Finally, our unexpected observation that in 3D HB and ICO cells form small and large diameter tubular structures that mirror the sizes of small peripheral biliary ductules and larger interlobular intrahepatic bile ducts, respectively, has important biological and clinical implications. The likelihood that lumen size is hard-wired will be an important consideration for liver tissue engineering where an optimal biliary network will form progressive connections between hepatocytic canaliculi, small ductules and larger ducts^32^. Moreover, the different tube sizes may model the poorly understood anatomical classes of human ICC and link that classification to the distinct epithelial states identified by multi-omics analysis, a possibility supported by our observation that lumen-competent FGFR2^fusion+^ ICC cell lines form small biliary ductule-sized lumens, consistent with their proposed classification into a ‘Bi-lineage’ epithelial state^43^. Our studies and model set the stage for dissecting the molecular basis of biliary lumen size and of the role of epithelial maturity in ICC tumorigenesis.

## Materials and Methods

### Cell culture

Single-cell-derived hepatoblast cell lines were generated from the livers of a litter of E14.5 mouse embryos as has been described^55,56^. This litter carried a homozygous unrecombined *Nf2^flox^* allele that has no detectable phenotypic consequences^57^. They were then cultured on collagen-coated plates in growth medium containing DMEM/F-12, 10% FBS (Avantor), 1% penicillin-streptomycin (P/S), 50ng/mL EGF (Peprotech), 30ng/mL IGFII (Peprotech), and 10μg/mL Insulin (Sigma). NIH3T3 cells were obtained from ATCC and grown on uncoated plates in DMEM, 10% calf serum, 1% P/S. HEK293T and Caco2 cells (ATCC) were grown on uncoated plates in DMEM, 10% FBS, 1% P/S. AML12 cells (ATCC) were grown on uncoated plates in DMEM/F12, 10% FBS, 1% P/S, 1% Insulin/Transferrin/Selenium liquid supplement (Sigma), 40ng/µL dexamethasone (Sigma). Human cholangiocytes from ICOs were established and maintained as previously described^33,44^. Briefly, ICOs were maintained in Matrigel domes in growth medium containing DMEM/F12, 1% P/S, 1% GlutaMAX, 10 mM HEPES, 1:50 B27 supplement (without vitamin A), 1:100 N2 supplement, 1 mM *N*-acetylcysteine, 10% (vol/vol) Rspo1-conditioned medium, 10 mM nicotinamide, 10 nM recombinant human [Leu^15^]-gastrin I, 50 ng/ml recombinant human EGF, 100 ng/ml recombinant human FGF10, 25 ng/ml recombinant human HGF, 10 μM Forskolin and 5 μM A83-01. A single cell suspension was generated by physically disrupting the basement membrane and digesting the organoids in TrypLE Express for seeding in 3D cell culture (described below). Patient derived biliary tract cancer cell lines (ICC10-6, ICC13-7, and ICC21) were generated as previously described^37^ and grown on uncoated plates in RPMI supplemented with 10% FBS and 1% P/S. All cells were cultured in a 37° C humidified incubator with 5% CO_2_. For examining receptor distribution by IF, cells were plated on collagen-coated glass coverslips (Corning) and grown for 3 days at 37° C prior to fixation.

For 3D cell culture, we modified a previously described method^31^. Collagen gel was prepared by mixing 80% type I collagen (Corning) by volume with 10% 10X PBS and 10% 0.1N NaOH. This was then mixed with growth factor-reduced, phenol red-free Matrigel (Corning 356231) in an 80:20 collagen gel-to-Matrigel ratio. 150μL of gel was layered into each well of a prechilled 8- well chamber slide with clear borders (Ibidi) and the gels were incubated for 2 h (hours) at 37° C, to solidify them. Between 1×10^5^ and 4 x 10^5^ cells, as indicated in the figure legends, were suspended in 450μL of growth medium plus 5ng/mL HGF (Peprotech), added to each well and grown for 2 days with 5% CO_2_ at 37° C. Medium was then removed and an additional 150 μL of gel of the same composition, followed by 300 μL of growth medium once the gels had been hardened for 2 h at 37° C, was layered on top. The cultures were incubated for an additional 3 days at 37° C prior to fixation. Due to the layout of 2 rows of 4 wells per slide, 3D experiments were done as technical duplicates. For phosphor-histone H3 (pHH3) quantifications, and where indicated in the figure legends, samples were instead fixed and stained without the addition of the second layer of gel to facilitate the antibody detection.

### Pharmacological inhibitors

For 3D experiments, compounds were added one day prior to gel overlay unless indicated otherwise in the figure legends at the following concentrations: Infigratinib (BGJ398; Selleckchem; 3 μM), PD98059 (Selleckchem; 1 μM), and dasatinib (Selleckchem; 1 μM). For immunoblotting, cells were starved for 20 h in medium without FBS or added growth factors, then switched to medium containing inhibitors for 3 h before harvesting. For qPCR, cells were treated with 3µM infigratinib for 16 h at steady state.

### Plasmids

The pCL-ECO, pCMV-VSVG, pMSCV-FGFR2^wt^, pMSCV-FGFR2-PHGDH and pMSCV-FGFR2-H167_N173 constructs were a generous gift from Nabeel Bardeesy^18,58^. The WI290>C substitution was generated in the pMSCV-FGFR2^wt^ construct using the Q5 site-directed mutagenesis kit (NEB). pMSCV-Blasticidin (empty vector) was a gift from David Mu (Addgene plasmid # 75085; http://n2t.net/addgene:75085 ; RRID:Addgene_75085). pUMVC was a gift from Robert Weinberg (Addgene plasmid #8449; http://n2t.net/addgene:8449 ; RRID:Addgene_8449).

### Antibodies

The following antibodies were used: E-cadherin (1:500 for IF, 1:2000 for immunoblot; BD Biosciences, clone 36), β-catenin (1:250 for IF; Abcam clone E247), Ezrin (1:500 for IF; Invitrogen clone 3C-12), N-cadherin (1:500 for IF, 1:2000 for immunoblot; BD Biosciences clone 32), phospho Histone H3 (1:200 for IF; Cell Signaling Technology #9701), FGFR2 (1:200 for IF, 1:1000 for immunoblot; Cell Signaling Technology, clone D4H9), phospho-FRS2 Y196 (1:1000 for immunoblot; Cell Signaling Technology #3864), total FRS2 (1:500 for immunoblot; R&D Systems clone 462910), β-tubulin (1:2000 for immunoblot; Sigma clone B-7), phospho-SHP-2 (1:1000 for immunoblot; Cell Signaling Technology #3751), phospho-MEK (1:1000 for immunoblot; Cell Signaling Technology #9121), total MEK (1:1000 for immunoblot; Cell Signaling Technology, #4694), phospho-Src family Y416 (1:1000 for immunoblot; Cell Signaling Technology clone D49G4), gm130 (1:500 for IF; BD Biosciences, clone 35), ZO-1 (1:100, Invitrogen #61-7300). HRP-conjugated secondary anti-mouse and anti-rabbit antibodies were from Amersham. Alexa Fluor 488, 555, and 647-conjugated secondary antibodies were from ThermoFisher and used at 1:500.

### Virus production and infection

Generation of MSCV retrovirus for infecting cells with FGFR2 constructs was achieved by PEI transfection of 1 μg pMSCV, 900 ng pCL-ECO (for mouse cells) or 900 ng pUMVC (for human cells) and 100 ng pCMV-VSVG per well of adherent HEK293T cells in a 6-well plate. Viral supernatants were collected at 2 and 3 days post-transfection. For infection, viral supernatants, containing 8 μg/mL polybrene, were added to cells grown to 30-50% confluence, then cells were selected and stable cell lines generated using blasticidin (Invivogen), at a concentration that had been experimentally determined to kill all uninfected cells after 7 days (12 µg/mL for HB^A^; 6 µg/mL for HB^B^; 7 µg/mL for HB^C^; 5 µg/mL for 3T3; 10 µg/mL for Caco2; 12 µg/mL for AML12).

### Immunofluorescence (IF)

For IF of 3D cultures, cells were fixed for 30 min (minutes) at 37° C with 3.7% formaldehyde in cytoskeletal buffer (10 mM MES Na^+^ Salt, 138 mM KCl, 3 mM MgCl_2_, 2 mM EGTA, pH 7.2), incubated for 15 min at room temperature with 100 U/mL collagenase (Sigma), permeabilized for 30 min at 37° C with 0.5% Triton in PBS (with Ca^++^ and Mg^++^ ions), and then washed 3X for 15 min with 100 mM glycine in PBS. Cells were then blocked for 1 h with 0.25% Triton in PBS, 0.1% BSA, 10% goat serum and incubated overnight at 4° C with primary antibodies. This was followed by incubations with Alexa Fluor-conjugated secondary antibodies and conjugated Phalloidin (1:200; ThermoFisher). Nuclei were stained with DAPI and samples were mounted with Prolong Gold (ThermoFisher).

For IF of cells on coverslips, cells were fixed in 3.7% PFA in PBS (with Ca^++^ and Mg^++^) for 15 min at room temperature, permeabilized with 0.2% Triton in PBS for 10 min, blocked for 30 min with 0.1% Triton in PBS, 0.1% BSA, 10% goat serum, and incubated overnight at 4° C with primary antibodies in blocking buffer. Cells were then incubated with Alexa Fluor-conjugated secondary antibodies, conjugated Phalloidin and DAPI.

Confocal images were captured with an LSM 710 or LSM 980 with Airyscan 2 inverted laser-scanning confocal microscope (Carl Zeiss) using 5x (EC Plan-Neofluar NA 0.16), 20x (Plan- Apochrmat NA 0.8), 40x multi-immersion (Plan-Apochromat NA1.2) or 63x oil-immersion (Plan-Apochromat NA 1.4) objectives (Carl Zeiss). DAPI was excited with the 405-nm diode laser.

Alexa Fluor 488, Alexa Fluor 555 and Alexa Fluor 647 probes were excited with the 445-nm diode laser. Images were acquired as single images or z-stacks in increments of 0.5 µm with the Airyscan 2 detector using ZEN Black software (2012; Carl Zeiss).

For HALO-mediated quantifications of lumens and dividing nuclei, samples were scanned using the Vectra 3.0 Automated Quantitative Pathology Imaging System, 6 slides (Vectra software version 3.05, Akoya). Using this widefield, multispectral imaging system, images were acquired at 20X magnification (NA 0.6). For each sample, the Vectra system captured the fluorescent spectra from 440nm to 720nm, using three filter cubes: AF488, AF647, and DAPI. These image captures were then combined to create a single stack image, which retained the unique spectral signature for each IF marker. Following an initial low power scan (4X), the region scanned was reviewed using Phenochart software, confirming the regions to be scanned using a higher power objective. Standard settings were used for multispectral image acquisition. For each scan, the focal plane was uniquely set to account for the 3D nature of the tissues. Multispectral deconvolution of multiplex IF stains was performed using inForm Image Analysis Software using a preloaded synthetic library (version 2.4.10, Akoya).

### qPCR

Total RNA was extracted from frozen cell pellets using TRIzol (Invitrogen), treated with DNase (NEB), then reverse transcribed with MMLV-RT (Promega) using oligo-dT primers. Fast Start Universal SYBR Green Mix (Millipore Sigma) was used to amplify 0.5 µL of the RT reaction in a 15 µL total reaction volume. Triplicate samples were run on a Light Cycler 480 system (Roche) with cycling conditions of denaturation for 15 seconds at 95° C, annealing for 1 min at 60° C, and extension at 60° C, 45 cycles. Expression was normalized to GAPDH and fold-change computed using the ΔΔCt method.

Primer sequences are as follows. *Cdhr5* forward 5′-TGCGCCAAAATTCTCCTTTGA-3′ and reverse 5′-AGGGATGACGGTTGTATTCACT-3′; *Epcam* forward 5′- GCGGCTCAGAGAGACTGTG-3′ and reverse 5′-CCAAGCATTTAGACGCCAGTTT-3′; *Gapdh*

forward 5′-AGGTCGGTGTGAACGGATTTG-3′ and reverse 5′-TGTAGACCATGTAGTTGAGGTCA-3′; *Hnf1b* forward 5′-CCCAGCAATCTCAGAACCTC-3′ and reverse 5′-AGGCTGCTAGCCACACTGTT-3′; *Sox9* forward 5′-ACTCTGGGCAAGCTCTGGAG-3′ and reverse 5′-CGAAGGGTCTCTTCTCGCTCT-3′.

### Western blotting

Cell pellets were lysed in mRIPA buffer with protease inhibitors (1% Triton X-100, 0.5% sodium deoxycholate, 0.1% SDS, 50 mM Tris-HCl at pH 7.4, 140 mM NaCl, 1 mM EDTA, 1 mM EGTA, 1 mM PMSF, 1 mM Na_3_VO_4_, 10 µg/mL leupeptin, 10 µg/mL pepstatin, 10 µg/mL aprotinin), for 1 h on ice, then lysates cleared by centrifugation. 40 µg of protein were loaded on SDS-PAGE gels, transferred to PVDF membranes, and incubated at 4° C overnight with antibodies diluted in 5% milk or 5% BSA (for phospho-specific antibodies). The membranes were then incubated with secondary antibodies, exposed to ECL substrate and film-developed.

### Enzymatic deglycosylation assay

To determine the glycosylation state, the cell lysates were treated with EndoH (New England Biolabs, P0702) or Peptide-N-Glycosidase F (New England Biolabs, P0704) as per manufacturer instructions. Briefly, the combination of 50 µg lysates and glycoprotein denaturing buffer was incubated at 100° C for 10 min. The denatured proteins were then treated with 2,000 U Endo H or 1,250 U PNGase F at 37°C for 1 h and analyzed by immunoblot.

### Cell surface biotinylation

Cells were seeded in 6-well plates, grown to confluence and then placed on ice for 10 min before rinsing 3X with ice-cold PBS containing Ca²⁺ and Mg²⁺. Biotinylation was performed by incubating the cells with 0.5 mg/mL EZ-Link Sulfo-NHS-SS-Biotin (ThermoFisher Scientific) in cold PBS at 4° C for 30 min with gentle swirling every 5 min to ensure even coating. After incubation, cells were rinsed 2X with cold PBS and washed 2X with cold 50 mM NH₄Cl in PBS to quench unbound biotin. Cells were then lysed in cytoskeleton (CSK) buffer (50 mM NaCl, 10 mM PIPES pH 6.8, 3 mM MgCl₂, 300 mM sucrose, 1% Triton-X 100, 10 mM NaF, 10 mM NaP₂O₇, 1X protease inhibitors) for 1 h on ice. Lysates were cleared by centrifugation at 14,000 x g for 30 min, and supernatants were incubated with streptavidin beads (ThermoFisher Scientific) for 2 h at 4° C with rotation. The beads were then washed 4X with CSK buffer, and biotinylated proteins eluted with 2x sample buffer containing β-mercaptoethanol at 95° C. Eluates and whole cell lysates as the input were analyzed by immunoblot.

### Image analysis and statistics

Confocal image analysis: ImageJ software (version 2.14, National Institutes of Health) was used for all confocal image processing and analysis. The displayed images were produced from single confocal slices or projections of z-stack images. Background was removed with rolling ball background subtraction. Lookup tables were applied to produce final images. x-z or y-z images were generated by re-slicing z-stack images. Cell height was quantified by measuring the length of a straight line drawn along E-cadherin labeled cell boundaries in x-z or y-z views of z-stack images. The junctional intensity of E-cadherin, N-cadherin or F-actin was determined by plotting the intensity profile of a line (10 μm in length) drawn perpendicular to the center of a cell-cell junction. The ratio of junctional to non-junctional intensity was calculated by dividing the average intensity of the middle of the line (∼3-7 μm) by the average intensity of the ends of the line (∼0-3 μm; 7-10 μm). Cord area was quantified by creating a binary threshold mask to define cords and measuring the percentage area covered by cords per field. Lumen polarity was measured in x-z views of z-stack images by drawing rectangles around the lumen perimeter as defined by F-actin staining and determining the Z coordinates of the center of the lumen relative to the center of a rectangle drawn around the full height of the cell monolayer. The absolute value of the difference between these values was graphed to represent the polarity of lumens relative to the monolayer.

Whole slide scanning analysis: After acquisition of whole-slide scans of 3D cultures, the spectrally unmixed images were imported into HALO^TM^ (Version 3.5) and fused. Annotation layers were selected to determine lumen and nuclear parameters within selected regions of the samples, typically the interior of the gel versus the exterior. Cell recognition and nuclear segmentation was trained and optimized using HiPlex FL module (Indica Labs), using both the DAPI and autofluorescence channels in at least five regions within each annotation. pHH3+ nuclei were counted in the AF647 channel, and the ratio of pHH3+ to total nuclei used as a measure of cell division. For lumen measurements, a supervised machine learning algorithm (Random Forest classifier) was trained on at least three randomly selected regions in the AF488 Phalloidin channel, to recognize lumens of differing sizes and shapes and to distinguish them from junctional Phalloidin. Data was output as number of lumens, lumen area, and lumen perimeter. For determining a lumen circularity index, the following formula was used: Circularity = (4π*Area)/(Perimeter^2^). Data from all analyses were imported into Prism 10 for plotting graphs and performing statistical analyses. The statistical tests used to compare groups are indicated in the figure legends.

### RNA sequencing sample preparation

Total RNA was extracted from frozen cell pellets using TRIzol reagent (Invitrogen) according to the manufacturer’s protocol. Residual DNA was removed by treatment with DNase I (New England Biolabs, M0303). The concentration of the extracted RNA was quantified using the Ribogreen fluorescence-based assay (Life Technologies) with a Victor X2 fluorometer (PerkinElmer). The integrity of the RNA was assessed using an Agilent Technologies 2100 Bioanalyzer, with samples achieving an RNA Integrity Number (RIN) value of 7.0 or higher considered suitable for further processing.

### RNAseq analysis

Libraries were prepared using the TruSeq Stranded mRNA LT Sample Prep Kit (Illumina) and sequenced on an Illumina NovaSeq6000 S4 v1.5 sequencer by Psomagen, which also provided processed quantification for each transcript as counts. RNAseqQC^59^ was used to assess library complexity, number of reads and genes detected, replicate variability and for hierarchical clustering of samples. Testing for differential gene expression between conditions was performed using DESeq2 (PMID: 25516281, v1.38.3). clusterProfiler (PMID: 34557778, v4.6.2) was used to perform gene set enrichment analysis (GSEA) with the Hallmark MSigDB genesets (PMID: 2677102, v7) and Gene Ontology over-representation analysis was done using the Cellular Components gene sets (PMID: 10802651, PMID: 36866529, 2022-07-01). We used the Benjamani-Hochberg method for multiple testing to adjust for FDRs (q). A p value <0.01 and a q value <0.01 were considered significant. Venn diagrams were generated from differential gene expression data and illustrate the overlap between genes that were upregulated in each condition (log_2_ fold change > 2, padj < 0.05). To compare the transcriptional profiles of our mouse HB cell lines with human ICC cell lines, mouse genes were first mapped to their human orthologs using the R biomaRt package (PMID: 19617889, PMID: 16082012, v2.62.1).

Following this, the R GSVA package (PMID: 23323831, v2.0.7) was used to calculate enrichment scores for each of the 4 ICC mRNA signatures in the mouse HB cell lines. P-values for differences in enrichment between signatures were calculated using the Mann-Whitney U test. All computational analyses were performed using custom scripts (available upon request) in R-4.2.2 or R-4.4.0.

## Acknowledgments

We thank past and present members of the McClatchey lab for valuable discussions, particularly Marcello Curto and Samira Benhamouche-Trouillet, and the Krantz Family Center for Cancer Research Cartography Core, which provided access to confocal microscopy equipment. We also thank Drs. Zhenghui G. Jiang and David Lee for providing materials used in deriving ICOs. This work was supported by the Harvard Stem Cell Institute (A.I.M.), Cholangiocarcinoma Foundation (E.O.), Children’s Tumor Foundation (E.O.), National Science Foundation (A.D.W.) and NIH (5R01DK127177-03, A.I.M.; 1R01DE033741-01A1, S.L.S; EB033821, S.N.B), and the MGH Research Scholars Fund (S.L.S). S.N.B. is a Howard Hughes Medical Institute Investigator.

## Abbreviations

CSK: Cytoskeleton buffer
EndoH: Endoglycosidase H
ERM: Ezrin, Radixin, and Moesin
EIDs: extracellular domain in-frame deletions
FGFR2: fibroblast growth factor receptor 2
FRS2: FGF receptor substrate 2
FGFRi: FGFR1-3 kinase inhibitor infigratinib
HB: hepatoblastCSK
IHBD: intrahepatic bile duct
ICC: intrahepatic cholangiocarcinoma
ICO: intrahepatic cholangiocyte organoids
MEKi: MAPK kinase inhibition
PNGase F: peptide-N-glycosidase F
pHH3: phospho-histone H3
PCA: principal component analysis
RTKs: receptor tyrosine kinases
SFKs: Src family tyrosine kinases

## References

1. Buckley, C.E. & St Johnston, D. Apical-basal polarity and the control of epithelial form and function. Nat Rev Mol Cell Biol 23, 559–577 (2022).

2. Whitford, M.K.M. & McCaffrey, L. Polarity in breast development and cancer. Curr Top Dev Biol 154, 245–283 (2023).

3. Peglion, F. & Etienne-Manneville, S. Cell polarity changes in cancer initiation and progression. J Cell Biol 223(2024).

4. Almagro, J., Messal, H.A., Elosegui-Artola, A., van Rheenen, J. & Behrens, A. Tissue architecture in tumor initiation and progression. Trends Cancer 8, 494–505 (2022).

5. Du, Z. & Lovly, C.M. Mechanisms of receptor tyrosine kinase activation in cancer. Mol Cancer 17, 58 (2018).

6. Clark, J.F. & Soriano, P.M. Pulling back the curtain: The hidden functions of receptor tyrosine kinases in development. Curr Top Dev Biol 149, 123–152 (2022).

7. Chiasson-MacKenzie, C. & McClatchey, A.I. Cell-Cell Contact and Receptor Tyrosine Kinase Signaling. Cold Spring Harb Perspect Biol (2017).

8. McDonell, L.M., Kernohan, K.D., Boycott, K.M. & Sawyer, S.L. Receptor tyrosine kinase mutations in developmental syndromes and cancer: two sides of the same coin. Hum Mol Genet 24, R60–66 (2015).

9. Lemaigre, F.P. Development of the Intrahepatic and Extrahepatic Biliary Tract: A Framework for Understanding Congenital Diseases. Annu Rev Pathol 15, 1–22 (2020).

10. Benhamouche-Trouillet, S., et al. Proliferation-independent role of NF2 (merlin) in limiting biliary morphogenesis. Development 145(2018).

11. Tanimizu, N., et al. Intrahepatic bile ducts are developed through formation of homogeneous continuous luminal network and its dynamic rearrangement in mice. Hepatology 64, 175–188 (2016).

12. Sigurbjornsdottir, S., Mathew, R. & Leptin, M. Molecular mechanisms of de novo lumen formation. Nat Rev Mol Cell Biol 15, 665–676 (2014).

13. Durdu, S., et al. Luminal signalling links cell communication to tissue architecture during organogenesis. Nature 515, 120–124 (2014).

14. Ryan, A.Q., Chan, C.J., Graner, F. & Hiiragi, T. Lumen Expansion Facilitates Epiblast-Primitive Endoderm Fate Specification during Mouse Blastocyst Formation. Dev Cell 51, 684–697 e684 (2019).

15. Fabris, L., et al. Pathobiology of inherited biliary diseases: a roadmap to understand acquired liver diseases. Nat Rev Gastroenterol Hepatol 16, 497–511 (2019).

16. Ilyas, S.I., et al. Cholangiocarcinoma - novel biological insights and therapeutic strategies. Nat Rev Clin Oncol 20, 470–486 (2023).

17. Vogel, A., Segatto, O., Stenzinger, A. & Saborowski, A. FGFR2 Inhibition in Cholangiocarcinoma. Annu Rev Med 74, 293–306 (2023).

18. Cleary, J.M., et al. FGFR2 Extracellular Domain In-Frame Deletions Are Therapeutically Targetable Genomic Alterations That Function as Oncogenic Drivers in Cholangiocarcinoma. Cancer Discov 11, 2488–2505 (2021).

19. Arai, Y., et al. Fibroblast growth factor receptor 2 tyrosine kinase fusions define a unique molecular subtype of cholangiocarcinoma. Hepatology 59, 1427–1434 (2014).

20. Wu, Y.M., et al. Identification of targetable FGFR gene fusions in diverse cancers. Cancer Discov 3, 636–647 (2013).

21. De Moerlooze, L., et al. An important role for the IIIb isoform of fibroblast growth factor receptor 2 (FGFR2) in mesenchymal-epithelial signalling during mouse organogenesis. Development 127, 483–492 (2000).

22. Chitnis, A.B., Nogare, D.D. & Matsuda, M. Building the posterior lateral line system in zebrafish. Dev Neurobiol 72, 234–255 (2012).

23. Antoniou, A., et al. Intrahepatic bile ducts develop according to a new mode of tubulogenesis regulated by the transcription factor SOX9. Gastroenterology 136, 2325–2333 (2009).

24. Zong, Y., et al. Notch signaling controls liver development by regulating biliary differentiation. Development 136, 1727–1739 (2009).

25. Loeuillard, E., Fischbach, S.R., Gores, G.J. & Ilyas, S.I. Animal models of cholangiocarcinoma. Biochim Biophys Acta Mol Basis Dis 1865, 982–992 (2019).

26. Kendre, G., et al. The Co-mutational Spectrum Determines the Therapeutic Response in Murine FGFR2 Fusion-Driven Cholangiocarcinoma. Hepatology 74, 1357–1370 (2021).

27. Sampaziotis, F., et al. Cholangiocyte organoids can repair bile ducts after transplantation in the human liver. Science 371, 839–846 (2021).

28. Babboni, S., et al. Cholangiocyte Organoids: The New Frontier in Regenerative Medicine for the Study and Treatment of Cholangiopathies. J Clin Med 13(2024).

29. Broutier, L., et al. Human primary liver cancer-derived organoid cultures for disease modeling and drug screening. Nat Med 23, 1424–1435 (2017).

30. Lidsky, M.E., et al. Leveraging patient derived models of FGFR2 fusion positive intrahepatic cholangiocarcinoma to identify synergistic therapies. NPJ Precis Oncol 6, 75 (2022).

31. Tanimizu, N., Miyajima, A. & Mostov, K.E. Liver progenitor cells fold up a cell monolayer into a double-layered structure during tubular morphogenesis. Mol Biol Cell 20, 2486–2494 (2009).

32. Jansen, P.L., et al. The ascending pathophysiology of cholestatic liver disease. Hepatology 65, 722–738 (2017).

33. Broutier, L., et al. Culture and establishment of self-renewing human and mouse adult liver and pancreas 3D organoids and their genetic manipulation. Nat Protoc 11, 1724–1743 (2016).

34. Valle, J.W., Lamarca, A., Goyal, L., Barriuso, J. & Zhu, A.X. New Horizons for Precision Medicine in Biliary Tract Cancers. Cancer Discov 7, 943–962 (2017).

35. Tanizaki, J., et al. Identification of Oncogenic and Drug-Sensitizing Mutations in the Extracellular Domain of FGFR2. Cancer Res 75, 3139–3146 (2015).

36. Pages, D.L., et al. Cell clusters adopt a collective amoeboid mode of migration in confined nonadhesive environments. Sci Adv 8, eabp8416 (2022).

37. Wu, Q., et al. EGFR Inhibition Potentiates FGFR Inhibitor Therapy and Overcomes Resistance in FGFR2 Fusion-Positive Cholangiocarcinoma. Cancer Discov 12, 1378–1395 (2022).

38. Lax, I., et al. The docking protein FRS2alpha controls a MAP kinase-mediated negative feedback mechanism for signaling by FGF receptors. Mol Cell 10, 709–719 (2002).

39. Clark, J.F. & Soriano, P. Diverse Fgfr1 signaling pathways and endocytic trafficking regulate mesoderm development. Genes Dev 38, 393–414 (2024).

40. Mitra, S.K. & Schlaepfer, D.D. Integrin-regulated FAK-Src signaling in normal and cancer cells. Curr Opin Cell Biol 18, 516–523 (2006).

41. Goeppert, B., Zen, Y., Valle, J., Klimstra, D. & Deshpande, V. Cholangiocarcinoma classification: current approach, relevance and challenges. J Clin Pathol 78, 298–299 (2025).

42. Nagtegaal, I.D., et al. The 2019 WHO classification of tumours of the digestive system. Histopathology 76, 182–188 (2020).

43. Vijay, V., et al. Generation of a biliary tract cancer cell line atlas identifies molecular subtypes and therapeutic targets. Cancer Discov (2025).

44. Westerfield, A.D., et al. A 3D <em>in vitro</em> model of the human hepatobiliary junction. bioRxiv, 2025.2007.2011.664464 (2025).

45. Dowbaj, A.M., et al. Mouse liver assembloids model periportal architecture and biliary fibrosis. Nature (2025).

46. Raab, M., et al. Van Gogh-like 2 is essential for the architectural patterning of the mammalian biliary tree. J Hepatol 81, 108–119 (2024).

47. Yanai, M., et al. FGF signaling segregates biliary cell-lineage from chick hepatoblasts cooperatively with BMP4 and ECM components in vitro. Dev Dyn 237, 1268–1283 (2008).

48. Azoury, S.C., Reddy, S., Shukla, V. & Deng, C.X. Fibroblast Growth Factor Receptor 2 (FGFR2) Mutation Related Syndromic Craniosynostosis. Int J Biol Sci 13, 1479–1488 (2017).

49. Bryant, D.M., Wylie, F.G. & Stow, J.L. Regulation of endocytosis, nuclear translocation, and signaling of fibroblast growth factor receptor 1 by E-cadherin. Mol Biol Cell 16, 14–23 (2005).

50. Nakamura, I.T., et al. Comprehensive functional evaluation of variants of fibroblast growth factor receptor genes in cancer. NPJ Precis Oncol 5, 66 (2021).

51. Chae, Y.K., et al. Phase II Study of AZD4547 in Patients With Tumors Harboring Aberrations in the FGFR Pathway: Results From the NCI-MATCH Trial (EAY131) Subprotocol W. J Clin Oncol 38, 2407–2417 (2020).

52. Nogova, L., et al. Evaluation of BGJ398, a Fibroblast Growth Factor Receptor 1-3 Kinase Inhibitor, in Patients With Advanced Solid Tumors Harboring Genetic Alterations in Fibroblast Growth Factor Receptors: Results of a Global Phase I, Dose-Escalation and Dose-Expansion Study. J Clin Oncol 35, 157–165 (2017).

53. Ruggieri, A.N., et al. Combined MEK/PD-L1 Inhibition Alters Peripheral Cytokines and Lymphocyte Populations Correlating with Improved Clinical Outcomes in Advanced Biliary Tract Cancer. Clin Cancer Res 28, 4336–4345 (2022).

54. Saha, S.K., et al. Isocitrate Dehydrogenase Mutations Confer Dasatinib Hypersensitivity and SRC Dependence in Intrahepatic Cholangiocarcinoma. Cancer Discov 6, 727–739 (2016).

55. Curto, M., Cole, B.K., Lallemand, D., Liu, C.H. & McClatchey, A.I. Contact-dependent inhibition of EGFR signaling by Nf2/Merlin. J Cell Biol 177, 893–903 (2007).

56. Strick-Marchand, H. & Weiss, M.C. Embryonic liver cells and permanent lines as models for hepatocyte and bile duct cell differentiation. Mech Dev 120, 89–98 (2003).

57. Giovannini, M., et al. Conditional biallelic Nf2 mutation in the mouse promotes manifestations of human neurofibromatosis type 2. Genes Dev 14, 1617–1630 (2000).

58. Goyal, L., et al. TAS-120 Overcomes Resistance to ATP-Competitive FGFR Inhibitors in Patients with FGFR2 Fusion-Positive Intrahepatic Cholangiocarcinoma. Cancer Discov 9, 1064–1079 (2019).

59. F, Z. R package (2024).

