## Supplemental Data for "Distinct phenotypic consequences of cholangiocarcinoma-associated FGFR2 alterations depend on biliary epithelial cell state"

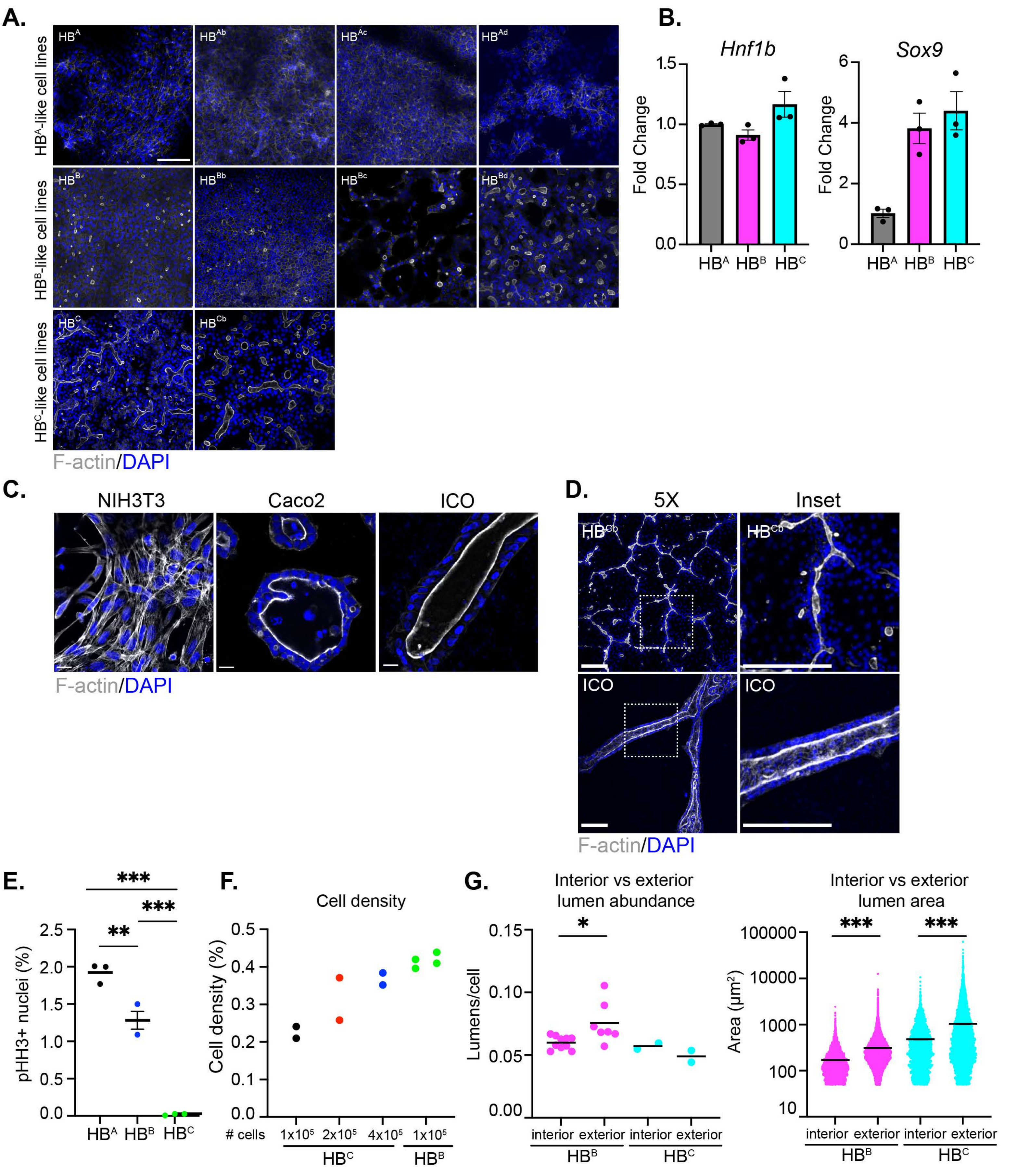
A.
HB^A-like cell lines
HB^B-like cell lines
HB^C-like cell lines
HB^A
HB^Ab
HB^Ac
HB^Ad
HB^B
HB^Bb
HB^Bc
HB^Bd
HB^C
HB^Cb
F-actin/DAPI
B.
Hnf1b
Sox9
Fold Change
1.5
1.0
0.5
0.0
6
4
2
0
HB^A HB^B HB^C
C.
NIH3T3
Caco2
ICO
F-actin/DAPI
D.
5X
Inset
HB^Cb
ICO
F-actin/DAPI
E.
pHH3+ nuclei (%)
2.5
2.0
1.5
1.0
0.5
0.0
***
**
***
HB^A HB^B HB^C
F.
Cell density
Cell density (%)
0.5
0.4
0.3
0.2
0.1
0.0
### cells
1x10^5 2x10^5 4x10^5 1x10^5
HB^C
HB^B
G.
Interior vs exterior lumen abundance
Lumens/cell
0.15
0.10
0.05
0.00
*
interior exterior interior exterior
HB^B
HB^C
Interior vs exterior lumen area
Area (µm²)
100000
10000
1000
100
10
***
***
interior exterior interior exterior
HB^B
HB^C

**A.**

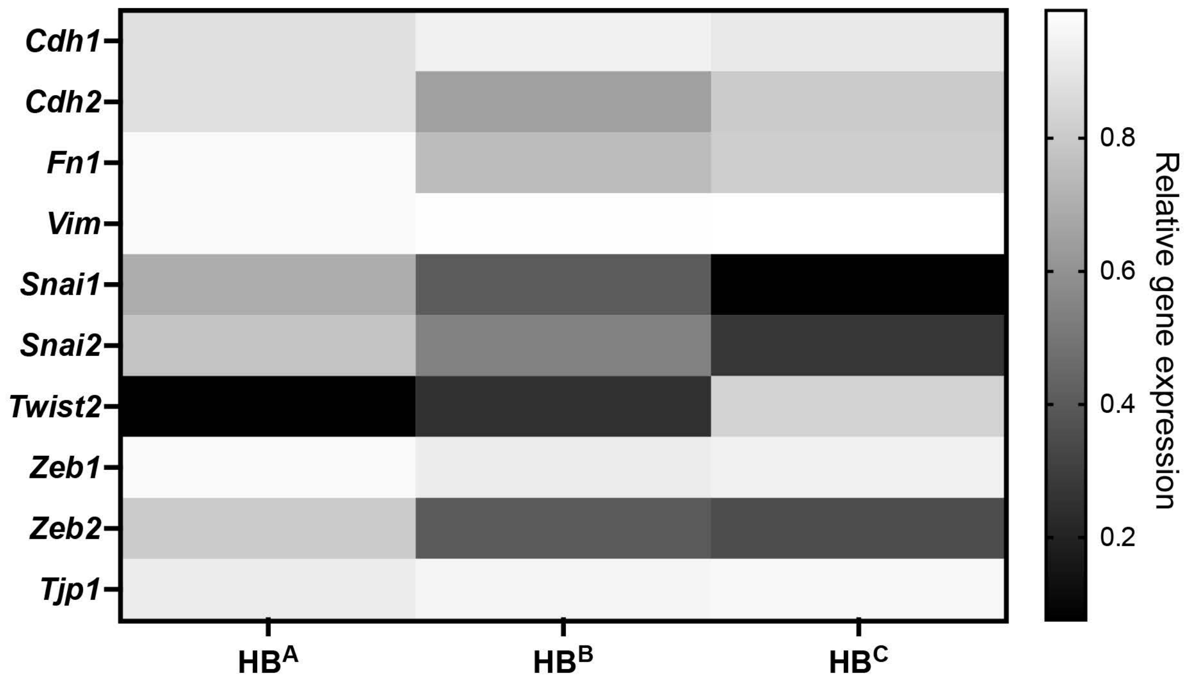

Cdh1
Cdh2
Fn1
Vim
Snai1
Snai2
Twist2
Zeb1
Zeb2
Tjp1
HB^A
HB^B
HB^C
0.8
0.6
0.4
0.2
Relative gene expression

Chiasson-MacKenzie, Zhang, O'Loughlin, et al. Supplemental Figure 3

**A.**

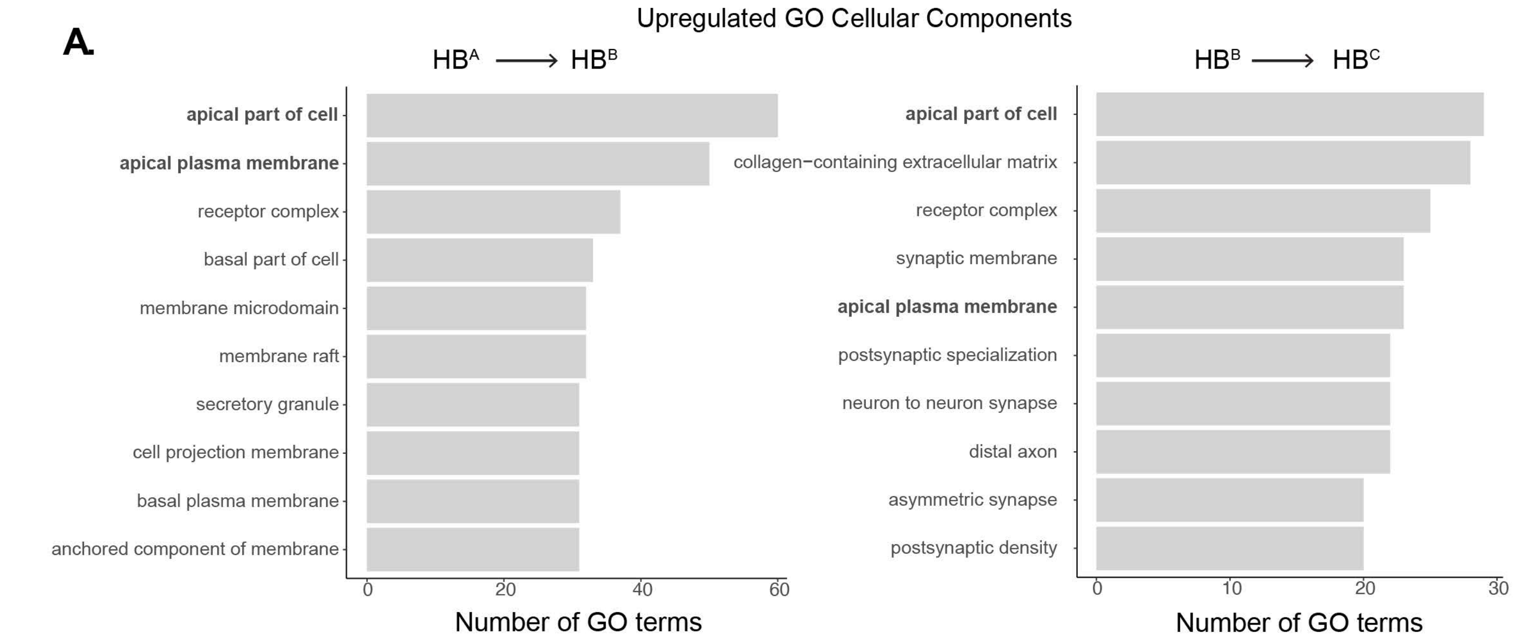


**B.**

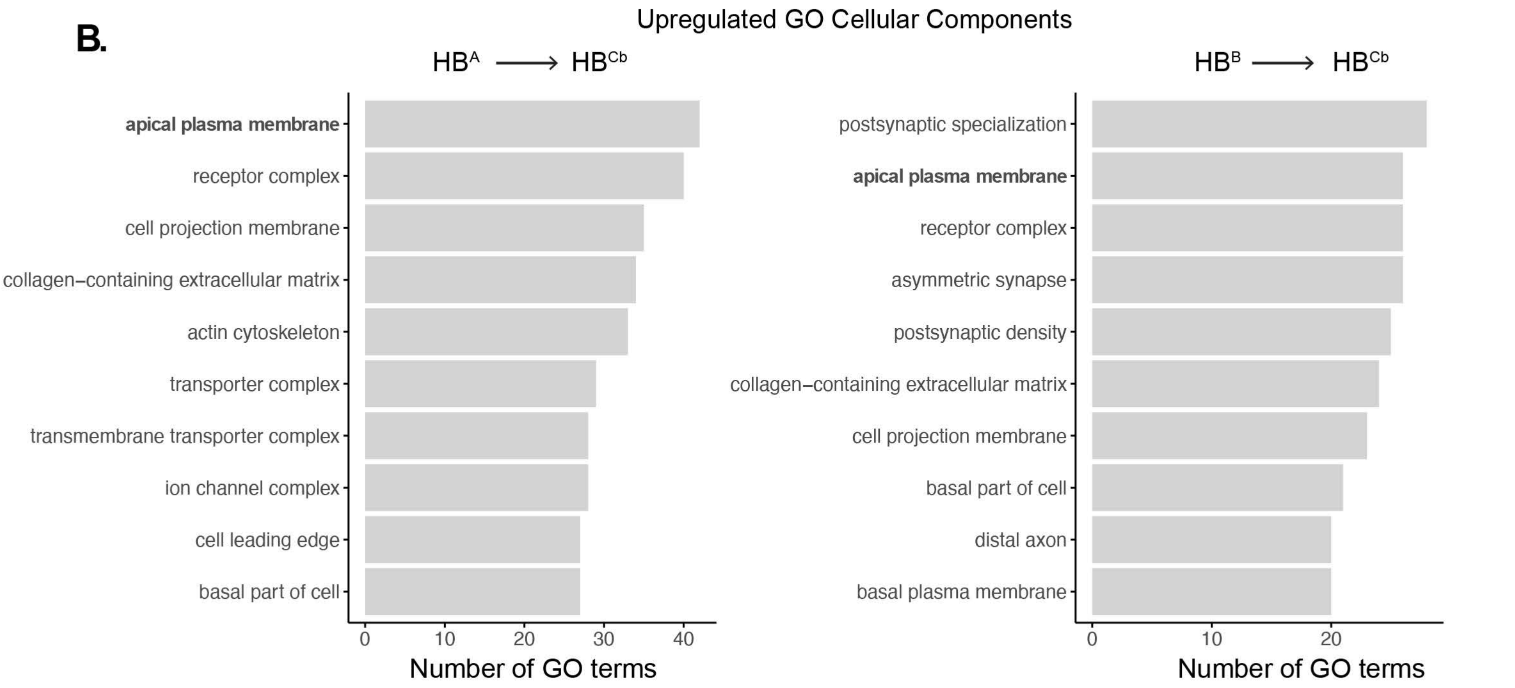

Chiasson-MacKenzie, Zhang, O'Loughlin, et al. Supplemental Figure 5

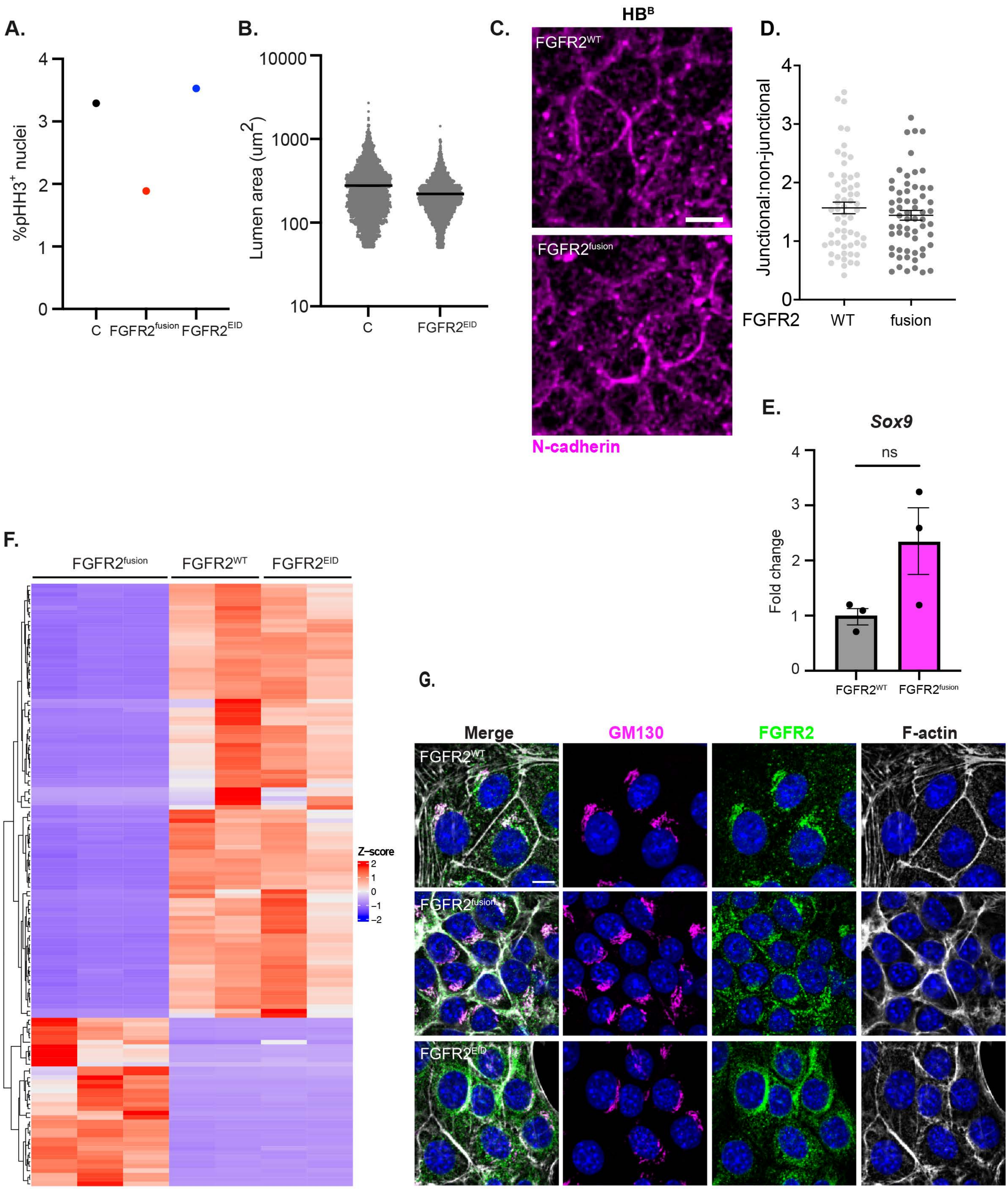

Chiasson-MacKenzie, Zhang, O'Loughlin, et al. Supplemental Figure 6

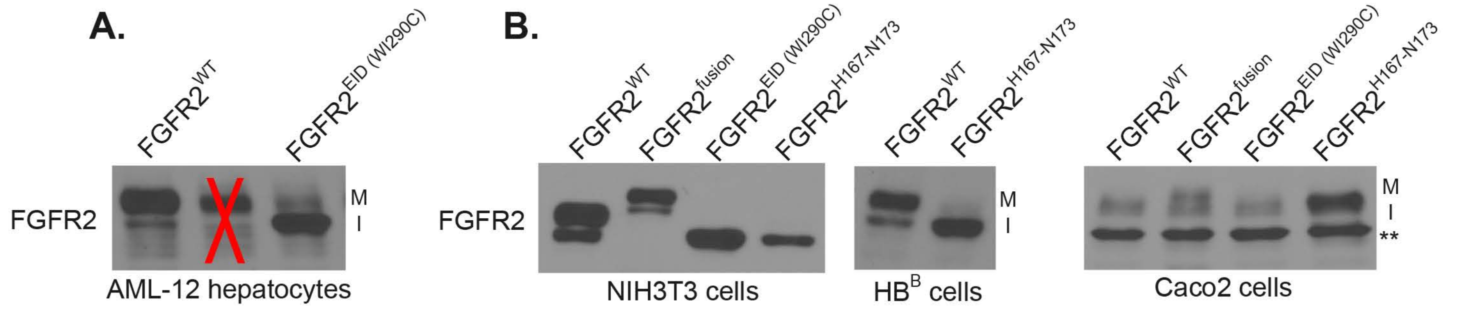

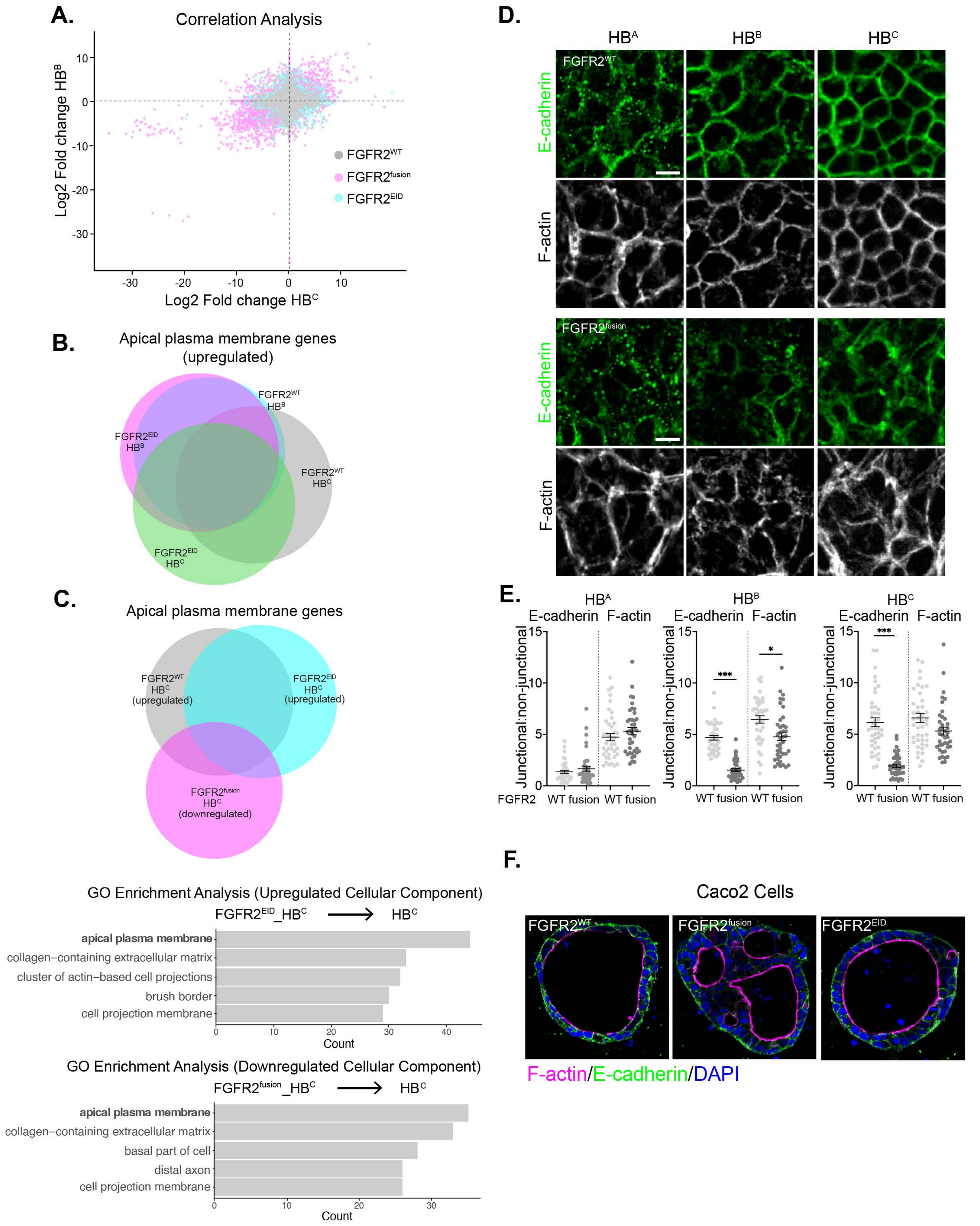
A.
Correlation Analysis
Log2 Fold change HB^B
Log2 Fold change HB^C
FGFR2^WT
FGFR2^fusion
FGFR2^EID
B.
Apical plasma membrane genes (upregulated)
FGFR2^WT HB^B
FGFR2^EID HB^B
FGFR2^WT HB^C
FGFR2^EID HB^C
C.
Apical plasma membrane genes
FGFR2^WT HB^C (upregulated)
FGFR2^EID HB^C (upregulated)
FGFR2^fusion HB^C (downregulated)
GO Enrichment Analysis (Upregulated Cellular Component)
FGFR2^EID_HB^C → HB^C
apical plasma membrane
collagen–containing extracellular matrix
cluster of actin–based cell projections
brush border
cell projection membrane
Count
GO Enrichment Analysis (Downregulated Cellular Component)
FGFR2^fusion_HB^C → HB^C
apical plasma membrane
collagen–containing extracellular matrix
basal part of cell
distal axon
cell projection membrane
Count
D.
HB^A
HB^B
HB^C
FGFR2^WT
FGFR2^fusion
E-cadherin
F-actin
E.
HB^A
HB^B
HB^C
E-cadherin F-actin
Junctional:non-junctional
FGFR2
WT fusion WT fusion
F.
Caco2 Cells
FGFR2^WT
FGFR2^fusion
FGFR2^EID
F-actin/E-cadherin/DAPI

Chiasson-MacKenzie, Zhang, O'Loughlin, et al.Supplemental Figure 8

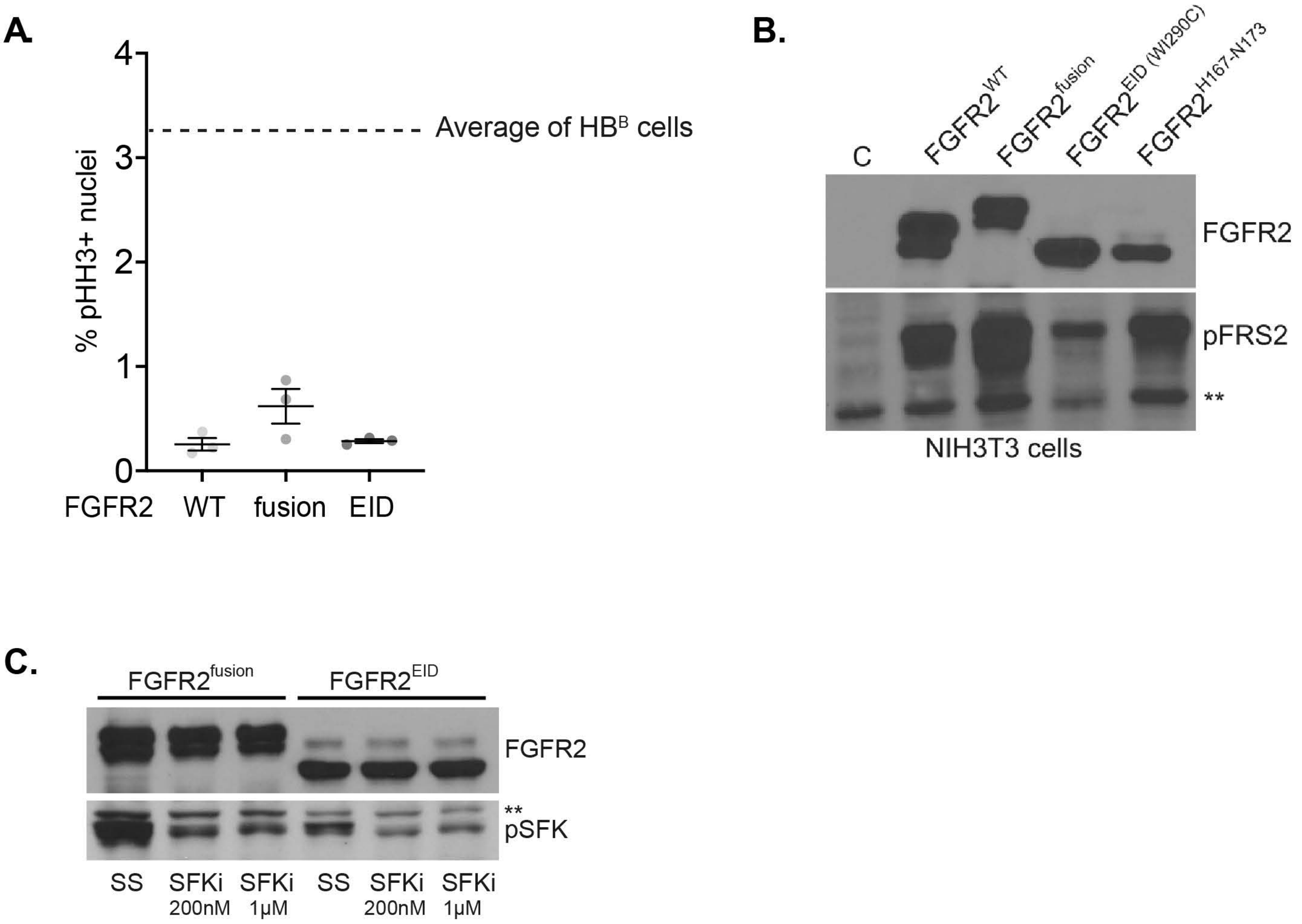

**A.**

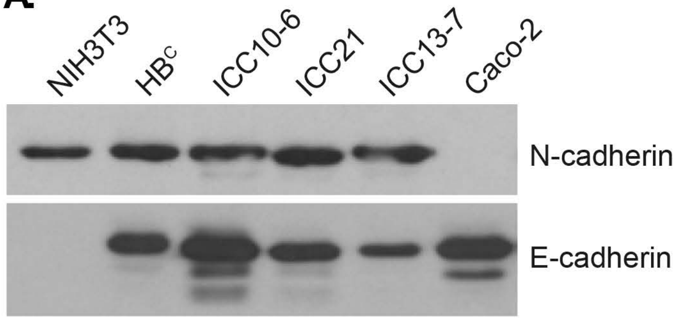

**Supplemental Figure Legends**

**Figure S1.** A. Representative confocal images showing F-actin (white) staining of lumens in all lumen-incompetent, immature lumen competent and mature lumen competent HB cell lines from our panel (blue, DAPI). Scale bar = 10 μm. B. Quantitation of *Hnf1b* and *Sox9* mRNAs in $HB^A$, $HB^B$ and $HB^C$ cells by qPCR. C. Confocal images showing F-actin (white) and DAPI (blue) staining of NIH3T3, Caco2, and human cholangiocyte ICO epithelial cells. Scale bars = 20 μm. D. Confocal images of the apical surfaces (F-actin, white) of tubular structures formed by HBC (small biliary ductule-like, 10-15 μm) and ICC (larger interlobular-like 30-60 μm) cells (blue = DAPI). Scale bars = 200 μm. E. Quantitation of pHH3+ nuclei in $HB^A$, $HB^B$ and $HB^C$ cells cultured on collagen/Matrigel without the overlay gel to facilitate staining. F. Quantitation of cell density in $HB^C$ and $HB^B$ cells seeded at different starting concentrations on collagen/Matrigel matrices. G. Quantitation of lumen abundance (left) and area (right; note log scale) in the interior (defined roughly as the area contained by a centrally placed rectangle of one half the dimensions of the full chamber) versus exterior (defined as the area outside of the rectangle) of individual chambers of $HB^B$ and $HB^C$ cells. Bars represents mean +/- SEM. P values were calculated using a one-way ANOVA with Tukey's multiple comparisons. (*$p<0.05$, **$p<0.01$, ***$p<0.001$).

**Figure S2**. Heatmap depicting the expression levels of various EMT markers from $HB^A$, $HB^B$ and $HB^C$ cells extracted from RNAseq data.

**Figure S3**. A. Bar graphs depicting top GO Cellular Components upregulated in $HB^B$ cells versus $HB^A$ cells and in $HB^C$ cells versus $HB^B$ cells. B. Bar Graphs depicting top GO Cellular Components upregulated in a second $HB^C$-like cell line ($HB^{Cb}$) versus $HB^B$ cells.

**Figure S5**. A. Quantitation of pHH3+ nuclei in control (C), FGFR2$^{fusion}$- and FGFR2$^{EID}$-expressing HB$^{B}$ cells. B. HALO-mediated quantification of lumen area in control (C) and FGFR2$^{EID}$-expressing HB$^{B}$ cells. C. Confocal images depicting the distribution of N-cadherin (magenta) in FGFR2$^{wt}$- or FGFR2$^{fusion}$-expressing HB$^{B}$ cells that were fixed without the addition of the overlay gel. D. Quantitation of junctional N-cadherin (magenta) in FGFR2$^{wt}$- and FGFR2$^{fusion}$-expressing HB$^{B}$ cells. E. Quantitation of Sox9 mRNA in FGFR2$^{wt-}$ versus FGFR2$^{fusion}$-expressing HB$^{B}$ cells by qPCR. F. Heatmap showing hierarchical clustering-based RNAseq profiles of HB$^{B}$ cells expressing FGFR2$^{fusion}$, FGFR2$^{wt}$ and FGFR2$^{EID}$. Grouped columns reflect RNAseq profiles of *independent passages* of each cell line as opposed to technical replicates. G. Confocal images of FGFR2$^{wt}$-, FGFR2$^{fusion}$- or FGFR2$^{EID}$-expressing HB$^{B}$ cells cultured on coverslips and stained for F-actin (white), GM130 (magenta) and FGFR (green). Scale Bars = 10 µm. Bars represents mean +/- SEM. P values were calculated using a two-tailed Student's t test (n.s. = not significant).

**Figure S6.** A. Immunoblot depicting FGFR2$^{wt}$ and FGFR2$^{EID}$ glycosylation patterns in murine AML12 hepatocytes. Red 'X' denotes an irrelevant lane. B. Immunoblots showing glycosylation patterns of FGFR2$^{wt}$, FGFR2$^{fusion}$ and two different FGFR2$^{EID}$s (WI290>C and H167_N173) expressed in NIH3T3 cells (left) and Caco2 cells (right). M = mature, I = immature, ** = irrelevant background band.

**Figure S7.** A. Correlation plot of FGFR2$^{wt}$, FGFR2$^{fusion}$, and FGFR2$^{EID}$ comparing Log2 fold change between HB$^{B}$ and HB$^{C}$ cells. B. Venn diagram showing the number of overlapping genes from the "apical plasma membrane" GO term (GO:0016324) whose expression is upregulated by FGFR2$^{WT}$ and FGFR2$^{EID}$ in HB$^{B}$ and HB$^{C}$ cells. C. (Top) Venn diagram depicting the number of overlapping genes from the "apical plasma membrane" GO term (GO:0016324) whose expression is upregulated by FGFR2$^{WT}$ and FGFR2$^{EID}$ and downregulated by FGFR2$^{fusion}$

in $HB^C$. (Bottom) Bar graphs depicting top GO Cellular Components upregulated in $FGFR2^{EID}$ $HB^C$ cells compared to control $HB^C$ cells and downregulated by $FGFR2^{fusion}$ in $HB^C$ compared to control $HB^C$ cells. D. Confocal images depicting the distribution of junctional E-cadherin (top, green) and F-actin (bottom, white) in $HB^A$, $HB^B$ and $HB^C$ cells expressing $FGFR2^{wt}$ (top) or $FGFR2^{fusion}$ (bottom). E. Quantitation of junctional E-cadherin or F-actin measured as the ratio of junctional to non-junctional intensity in $FGFR2^{wt}$ or $FGFR2^{fusion}$-expressing $HB^A$ (left), $HB^B$ (middle) and $HB^C$ (right) cells. P values were calculated using a one-way ANOVA with Tukey's multiple comparisons test (*p<0.2, ***p<0.001). F. Confocal images of Caco2 cells expressing $FGFR2^{wt}$, $FGFR2^{fusion}$, or $FGFR2^{EID}$ and stained for F-actin (magenta), E-cadherin (green) and DAPI (blue). Scale bars = 10 μm.

**Figure S8.** A. HALO-mediated quantitation of pHH3+ nuclei from $FGFR2^{wt}$-, $FGFR2^{fusion}$- and $FGFR2^{EID}$-expressing $HB^C$ cells. Dotted line represents the percentage of pHH3+ $HB^B$ cells expressing $FGFR2^{wt}$ for comparison (see Fig. S5A). Bars represents mean +/- SEM. B. Immunoblot showing the levels of pFRS2 triggered by $FGFR2^{wt}$, $FGFR2^{fusion}$ and two different $FGFR2^{EID}$s (WI290>C and H167_N173) expressed in NIH3T3 cells. Note that the top blot is the same as Fig S6B.

**Figure S9.** A. Immunoblot showing the levels of E-cadherin and N-cadherin protein across ICC10-6, ICC13-7, and ICC21 tumor cell lines along with controls of NIH3T3 (N-cadherin only), $HB^C$ (N- and E-cadherin), and Caco-2 cells (E-cadherin only).
